# Localized immunomodulation with cytokine-producing cells to mitigate host immune rejection responses in rodents and a non-human primate

**DOI:** 10.1101/2025.10.13.682221

**Authors:** Boram Kim, Dilrasbonu Vohidova, Amanda Nash, Yuen San Chan, Samantha Fleury, Shravani Deo, Danna Murungi, Peter D. Rios, Ira Joshi, Hafsa Nasir, Daisy Lopez, Mor Sela Golan, Cassidy Hart, Jose Oberholzer, H. Courtney Hodges, Omid Veiseh

## Abstract

The efficacy of cell-based therapeutics is often compromised by host immune recognition of implanted cells and biomaterials, resulting in fibrotic encapsulation and loss of function. Here, we address this challenge with an immunomodulatory cell-based therapy, in which alginate-encapsulated retinal pigment epithelial cells continuously secrete cytokines to locally modulate the implant microenvironment. In a healthy rodent model, the localized production of interleukin-10 (IL-10) or IL-12 from encapsulated cytokine-producing cells prevented host immune rejection and fibrosis of alginate capsules. Mechanistically, treatment was associated with reduced expression of pro-fibrotic genes and immune shifts consistent with macrophage and T-cell regulation, supporting a cytokine-mediated mitigation of foreign body response. In a diabetic murine model (streptozotocin-induced C57BL/6J), co-implantation of human islets with IL-10-producing cells attenuated pericapsular fibrosis, preserved islet viability, and restored normoglycemia for up to 100 days (4.76 times longer than islets alone). Significantly, IL-10-producing cells were also effective in enabling the durability and function of encapsulated cells in a healthy non-human primate, showing translational feasibility. Collectively, these findings suggest that localized cytokine delivery can reduce fibrotic encapsulation and support durable graft function, offering a path to lessen reliance on systemic immunosuppression in islets transplantation and other implantable biomaterial therapies.

**Teaser:** Encapsulated IL-10-producing cells locally suppress fibrosis and extend graft function in rodent models and a non-human primate.

## INTRODUCTION

Biomaterials are at the forefront of advances in regenerative medicine with applications in medical implants, surgical devices, drug delivery systems, cell-based therapeutics, wound healing, and more. However, one of the biggest remaining challenges for biomaterials is overcoming the foreign body response (FBR). The FBR is a naturally occurring immunological reaction to the introduction of foreign, or “non-self” medical implants, devices, or biomaterials (*1-3*). This inflammatory response is orchestrated primarily by macrophages and fibroblasts and results in the cellular deposition of a dense fibrous connective tissue layer on and around the foreign material (*3, 4*). The FBR ultimately leads to isolation of the medical device or biomaterial from the surrounding tissue and subsequent loss of function (*2*). Recent studies have demonstrated that the surface properties (e.g., surface chemistry, charge, topography, material composition, and wettability) of the implanted material play a critical role in delaying, dampening, or circumventing the foreign body reaction (*1, 5-9*).

Immune recognition of foreign materials initiates a cascade of cellular processes including inflammation, the formation of foreign body giant cells (fused accumulation of macrophages), and fibrosis (cellular deposition of collagenous coating to the surface of the foreign material) (*3*). The major cells involved in the recognition of foreign materials are macrophages which use pattern recognition surface receptors to separate identification of “self” from “non-self” (*10, 11*). If the foreign material is small enough, macrophages will attempt to directly eliminate the material via phagocytosis. However, if the foreign material is too large for the macrophage to engulf, which is often the case for implantable devices and drug delivery systems, the macrophages will instead accumulate and fuse to form multinucleated foreign body giant cells (FBGC) to attempt to degrade or eliminate the material (*3*). Other innate immune cells, such as neutrophils, will migrate to the area with the foreign material to help fight off any potential infections associated with the intrusion (*12*). If even the FBGCs cannot eliminate the foreign material, the macrophages will signal for fibroblasts to initiate fibrosis of the impervious material. Fibrosis is the accumulation (or in this case, deposition) of excess dense extracellular matrix components (such as collagen) on the surface of the foreign material to encapsulate and isolate it from the surrounding cells and tissues (*9, 13-16*). The deposition of fibrotic overgrowth is what ultimately leads to loss of function for many medical devices and implants. For this reason, scientists have been carefully studying biomaterials, surface properties, and other immunomodulatory techniques, to identify the optimal methods and materials to circumvent (or delay for as long as possible) the naturally occurring FBR.

Because macrophages orchestrate these events through cytokine signaling, tuning the local cytokine environment is a rational way to modulate FBR. Within this cytokine network, interleukin-10 (IL-10) limits macrophage activation, promotes regulatory T cells (T_reg_), and has been shown in preclinical implant models to attenuate fibrosis (*17*). By contrast, IL-12, while classically pro-inflammatory through Th1/IFN-γ pathways, can exert tissue- and context-dependent anti-fibrotic effects via IFN-γ-mediated antagonism of TGF-β signaling (*18, 19*). To date, most cytokine therapies in preclinical and clinical settings have been delivered systemically (e.g., intravenous/subcutaneous dosing or systemic depots), approaches that are often constrained by short half-life, peak systemic exposure, and the need for repeated dosing (*20-22*). These practical limitations motivate compartmentalized, local delivery to recalibrate the immune environment at the graft interface while minimizing systemic immunosuppression.

Localized immunomodulation via the administration of exogenous cytokines has been highlighted recently as a promising method of altering the immunological landscape or response/function within an isolated cavity, such as the abdominal cavity. We employ local cell delivery using alginate capsules, a well-studied carrier with mild gelation, biocompatibility, and tunable permeability (*3, 9, 15, 16, 23*). Despite physical barriers restricting direct cellular contact between host and donor cells, alginate capsules are permeable to secreted factors, which can be recognized as foreign antigens by the recipient’s immune system, triggering an alloreactive immune response (*24, 25*). This may trigger the FBR, ultimately leading to fibrosis and graft failure (Fig. 1A) (*26-31*). Building on our prior work, we previously validated an alginate-encapsulated retinal pigment epithelial (RPE) “cytokine factory” platform in which RPE cells secreted IL-2 for local cancer immunotherapy (*14, 15*). While IL-2 itself is not anti-fibrotic and that application targeted T-cell activation, those studies established the platform’s biocompatibility, manufacturability, containment, and capacity for sustained in vivo cytokine release. Here, we repurpose the same modular delivery system to release anti-fibrotic cytokines for FBR mitigation in the context of islet transplantation. Such localized delivery strategies may be particularly valuable for improving the clinical utility of islet transplantation, especially in light of recent advances such as the FDA approval of Lantidra (*32, 33*) and Vertex’s ongoing Phase 3 trials of stem cell-derived islets (NCT04786262, NCT06832410), both of which currently rely on systemic immunosuppression. In contrast, local immune modulation enables precise control over the immune response at the graft site, reducing systemic toxicity while enhancing graft protection and long-term function.

**Fig. 1.**
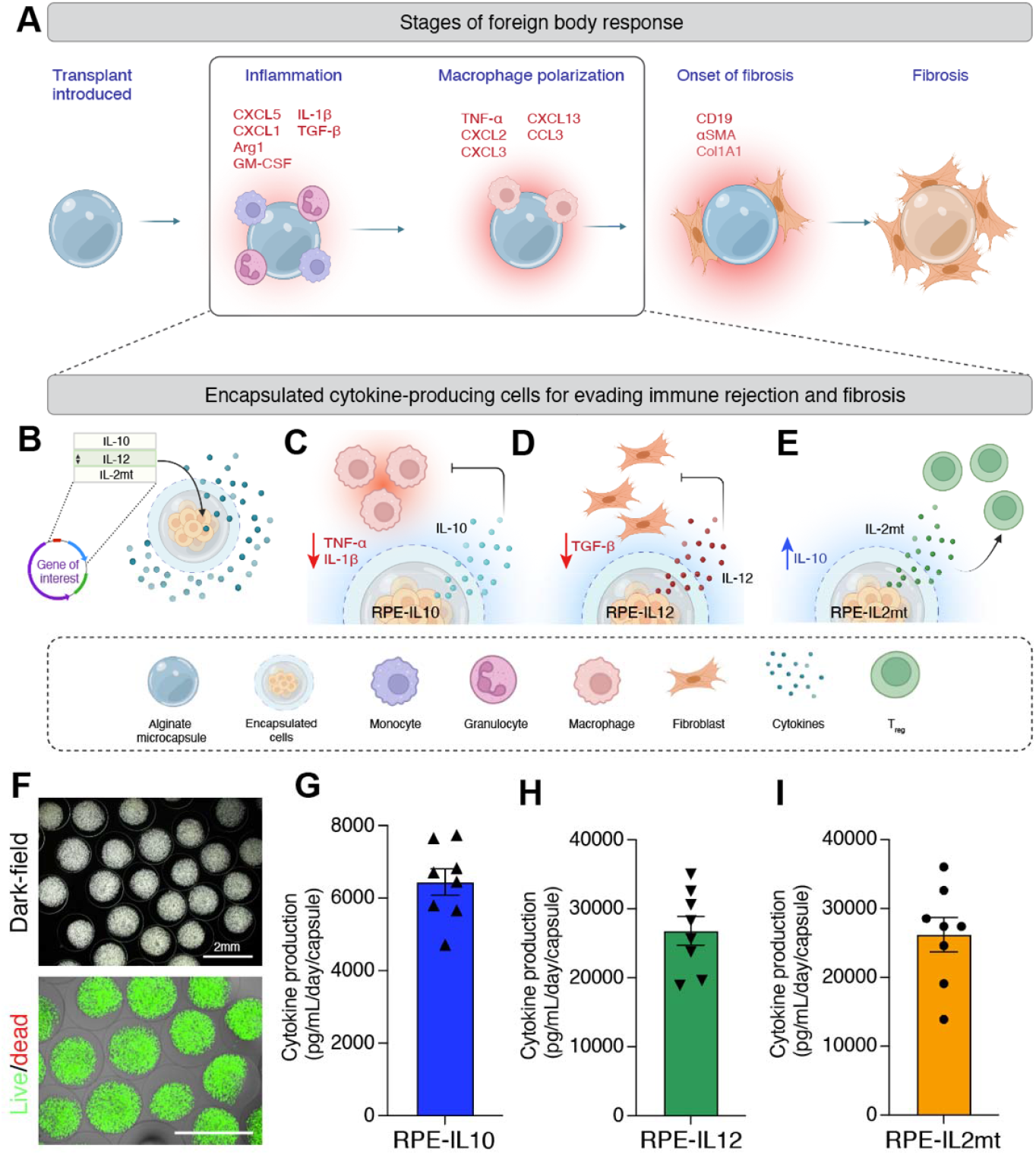
Cytokines released from encapsulated engineered cells modulate the foreign body response. (**A**) Schematic of fibrosis progression in alginate microcapsules, illustrating key immune cells, cytokines, and chemokines involved in the foreign body response (FBR), based on previously reported data (*3, 38*). FBR is initiated by protein adsorption, triggering early inflammation, followed by infiltration of granulocytes and monocytes at the implant site. Granulocytes express elevated levels of CXCL5 and IL-1β, while monocyte-derived macrophages secrete granulocyte-macrophage colony-stimulating factor (GM-CSF), arginase-1 (Arg1), transforming growth factor-β (TGF-β), and CXCL1, which contribute to fibrotic signaling. As fibrosis progresses, TGF-β expression increases, alongside chemokines such as CXCL2, CXCL3, CXCL13, and CCL3 that promote immune cell recruitment and macrophage polarization. In an attempt to phagocytose the implant, macrophages fuse into foreign-body giant cells (FBGCs). High local TGF-β levels drive FBGC formation and the conversion of fibroblasts into myofibroblasts, marked by the increased presence of α-smooth muscle actin (αSMA) and collagen 1A1 (Col1A1)-positive cells over time, indicating the onset of fibrosis. (**B**) Human retinal pigment epithelial (RPE) cells engineered to locally secrete therapeutic cytokines. (**C**) IL-10 suppresses macrophage activation by inhibiting pro-inflammatory cytokine production. (**D**) IL-12 promotes IFN-γ production, which downregulates TGF-β and restricts myofibroblast differentiation. (**E**) IL-2mt selectively expands IL-10-producing regulatory T cells (T_reg_) due to its reduced affinity for the IL-2Rβγ receptor. (**F**) Dark-field (top) and live/dead fluorescence (bottom) images of encapsulated cytokine-producing cells; live cells appear green, and dead cells appear red. Scale bar, 2 mm. (**G** to **I)**, Cytokine secretion levels from encapsulated engineered RPE cells over 24 hours measured via ELISA: RPE-IL10 (G), RPE-IL12 (H), and RPE-IL2mt (I).

In this study, we developed an immunomodulatory cell platform, adapted from our validated IL-2 “cytokine factory” system (*14, 15*), that uses engineered RPE cells to constitutively secrete immunomodulatory cytokines into the local microenvironment. Encapsulation in alginate protects the cells from direct host interaction, while enabling precise cytokine delivery for effective immune modulation. We selected human RPE cells due to their favorable clinical experience, low immunogenicity, and high secretory capacity (*14, 15, 34*). Specifically, we engineered RPEs to secrete IL-10, IL-12, or an IL-2 mutein (IL-2mt) (Fig. 1B). IL-10 limits macrophage activation and dampens pro-inflammatory mediators such as TNF-α and IL-1β (Fig. 1C) (*17*); IL-12 can counter-regulate profibrotic TGF-β programs via IFN-γ (Fig. 1D) (*18, 19*); and IL-2 mutein (IL-2mt) preferentially expands T_reg_ through reduced IL-2Rβγ engagement, which can inhibit fibroblast proliferation and potentially prevent or delay fibrosis (Fig. 1E) (*35-37*). We evaluated this platform for pericapsular fibrosis control around encapsulated islets in streptozotocin (STZ)-induced C57BL/6J mice, and assessed translational feasibility in a pilot non-human primate (NHP) study with safety and pharmacokinetic monitoring of localized IL-10 delivery. Together, these studies test whether localized cytokine production can mitigate fibrosis and support durable graft function while reducing reliance on systemic immunosuppression. This localized immunomodulatory approach presents a clinically translatable strategy not only to improve the durability and safety of islet transplantation, but also to broadly address immune-mediated fibrosis and rejection in implanted biomaterials, and potentially to treat other autoimmune diseases through targeted, site-specific immune regulation.

## RESULTS

### Fabrication of cytokine-producing cells

To evaluate the efficacy of cytokine-producing RPE cells in preventing biomaterial-induced fibrosis, three distinct cytokines with immune-modulating potential (e.g., IL-10, IL-12, and IL-2mt) were selected. RPE cells underwent genetic modification to express a specific immunomodulatory cytokine through the PiggyBac transposon system. This system enables swift prototype development by substituting the gene of interest while preserving the optimized backbone. We refer to encapsulated cytokine-producing RPE cells as RPE-IL10, RPE-IL12, or RPE-IL2mt, depending on the specific cytokine they were engineered to produce. The engineered cells were encapsulated, as previously described (*14, 15*), in 1.5mm alginate capsules containing ∼10,000 cells per capsule, showing highly viable cells after encapsulation (Fig. 1F). Following encapsulation, cytokine secretion from preimplant capsules was confirmed using ELISA after 24 hours of incubation (Fig. 1, G to I). RPE-IL10 exhibited a production rate of approximately 6.4 ng/mL/day/capsule, RPE-IL12 yielded about 25.7 ng/mL/day/capsule, and RPE-IL2mt produced about 26.2 ng/mL/day/capsule.

### Local delivery of anti- or pro-inflammatory cytokines can prevent fibrosis on the surface of biomaterials

To evaluate the anti-fibrotic effects of cytokine-producing capsules, a mix of large (1.5 mm) and small (0.5 mm) capsules were implanted in the intraperitoneal (IP) space of healthy C57BL/6J mice. Large capsules contained cytokine-producing cells, delivering a controlled, low-dose treatment (100,000 cells per mouse) in a minimal volume. The 0.5 mm microcapsules, known to trigger immune responses (*3*), were included to create a pro-fibrotic environment while remaining suitable for future islet encapsulation studies. Each mouse received 10 large capsules containing cytokine-producing cells (1.5 mm) along with ∼400 µL of empty microcapsules (0.5 mm) in IP space (Fig. 2A). This design ensured cytokine delivery in a minimal implant volume while allowing assessment of anti-fibrotic effects against a robust immune response.

**Fig. 2.**
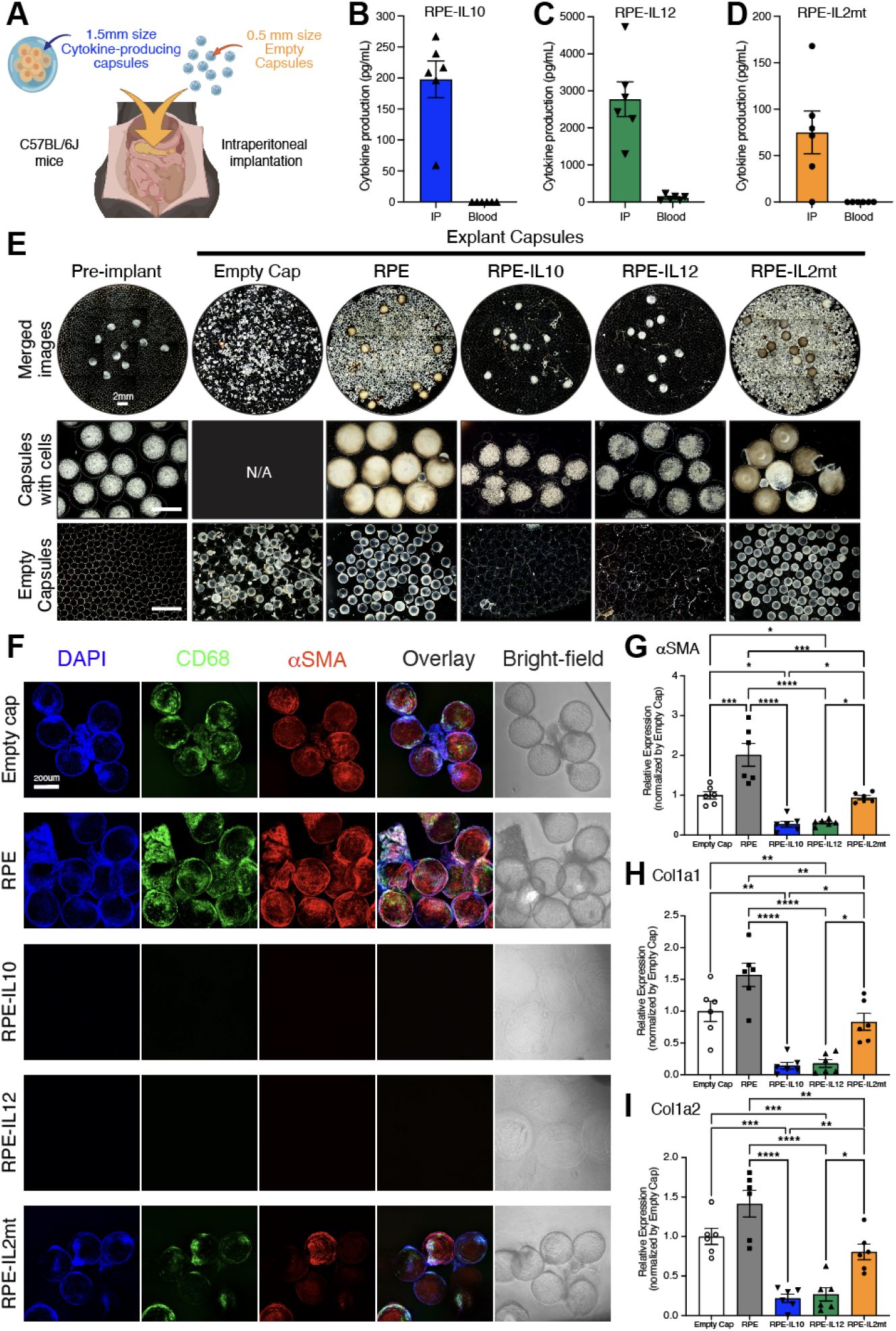
Co-delivery of cytokine-producing capsules can prevent fibrosis of biomaterials in the short term. (**A**) Schematic of local cytokine delivery in IP space of C57BL/6J mouse for one month. (**B** to **D**) Cytokine productions in local (IP) and systemic (blood) levels after 1month implantations of RPE-IL10 (B), RPE-IL12 (C), and RPE-IL2mt (D) groups, respectively (n = 6). (**E**) Representative dark-field images of pre-implant and explante capsules. Scare bar, 2mm. (**F**) Immunofluorescence images of explanted microcapsules. DAPI, blue; CD68, green; αSMA, red. Scale bar, 200 μm. (**G** to **I**) RT-qPCR analysis to compare fibrotic gene expressions. αSMA (G), Col1a (H), and Col1a2 (I) normalized to empty capsule control. All graph bars are mean ± s.e.m of biological replicates (n = 6). Two-way ANOVA with Bonferroni correction was used for statistical analysis (****p < 0.0001, ***p < 0.0002, **p < 0.002, *p < 0.033).

First, the concentration of cytokines in the local (IP space) and systemic cavity (blood) was assessed one month after implantation to determine the biodistribution of our cytokine-producing capsules (Fig. 2, B to D). Minimal levels of cytokines were detected in the bloodstream for all three cytokine-producing capsules suggesting that the administered cytokines were contained within the local cavity. RPE-IL10 (Fig. 2B) and RPE-IL12 (Fig. 2C) exhibited high local concentrations in IP fluids (∼195pg/mL and ∼2625pg/mL, respectively), indicating sustained cytokine production in the local space post-1-month implantation. RPE-IL2mt (Fig. 2D) did not demonstrate high local concentration (∼75.5pg/mL). Subsequently, all microcapsules were retrieved and analyzed with microscopy imaging for fibrosis deposition on their surfaces (Fig. 2E, fig. S1). The groups containing non-engineered RPE or RPE-IL2mt displayed highly packed fibrotic overgrowth on surface of the capsules in all mice, as evidenced by dark-field images. In contrast, the RPE-IL10 and RPE-IL12 groups showed minimal overgrowth on the 1.5mm cell containing capsules and the empty microcapsules suggesting that the presence of IL-10 or IL-12 played a role in mitigating the fibrotic process. The empty capsule group still exhibited fibrotic responses in the absence of xenogenic cells, consistent with previous reports indicating that smaller capsule sizes can elicit more robust immune responses (*3*). Explanted RPE-containing capsules were stained with a live/dead assay (live, green; dead, red) to assess viability (fig. S1). In the RPE and RPE-IL2mt groups, capsules were densely encased by pericapsular tissue; fluorescence was largely confined to cells on the capsule surface, and the encapsulated RPEs were obscured. By contrast, IL-10 and IL-12 groups showed little to no surface fibrosis, and viable RPEs were readily visualized within the capsules, with strong live (green) signal. In addition, a high-dose condition was evaluated by loading engineered RPE cells at 40,000 cells per capsule (total 400,000 cells per mouse). RPE, RPE-IL10, and RPE-IL2mt groups were compared one month post-implantation, with 0.5 mm microcapsules co-delivered as in the low-dose condition (10,000 cells per capsule). High-dose IL-12 was not included due to dose-limiting cytotoxicity (*34*). Under high-dose conditions, RPE-IL10 again exhibited effective fibrosis prevention, showing minimal overgrowth on explanted capsule surfaces relative to RPE controls (fig. S2). RPE-IL2mt did not achieve the level of fibrosis prevention observed with IL-10 despite higher cytokine production. On this basis, subsequent studies were conducted under the low-dose regimen to minimize implanted cell burden while retaining anti-fibrotic efficacy.

Next, immunofluorescent imaging was utilized to evaluate the extent of fibrotic overgrowth and cellular deposition on the surface of the explanted capsules (Fig. 2F). Retrieved microcapsules were stained with alpha-smooth muscle actin (αSMA), a myofibroblast marker, and CD68, a macrophage marker. Results were consistent with dark-field imaging, revealing highly visible fibrotic marker expression on the capsule surface of the empty cap, RPE, and RPE-IL2mt groups but not RPE-IL10 and RPE-IL12 groups. Finally, RT-qPCR was conducted to confirm the expression of fibrosis-related genes using cells collected from retrieved microcapsules (Fig. 2, G to I). The relative expression of αSMA (Fig. 2G), Col1a1 (Fig. 2H), and Col1a2 (Fig. 2I) was compared among the groups, and the results aligned with the microscopic findings. These results suggest that co-delivery of either IL-10 or IL-12-producing capsules significantly reduced fibrosis on empty microcapsules compared to the non-engineered cell delivery group.

To understand the medium and long-term anti-fibrotic effect of RPE-IL10 and RPE-IL12, we conducted a similar in vivo experiment to the experiment described above. Briefly, 1.5mm RPE-IL10 or RPE-IL12 capsules were co-administered with empty micro-capsules within the IP space of healthy C57BL/6J mice for 3 months or 6 months. At each timepoint, capsules were retrieved and evaluated under the microscope as described above. All capsules explanted from the RPE-IL10 group demonstrated minimal fibrotic overgrowth on their surface, suggesting sustained prevention of fibrosis (fig. S3). Only few mice exhibited partial fibrosis deposition on the surface of the IL-10 capsules after three months (fig. S3A) and six months (fig. S3B). Similar outcomes were observed in the RPE-IL12 group (fig. S4), indicating robust anti-fibrotic immune modulation (with the exception one mouse out of six from the 6-month post-implantation group). To assess biocompatibility and downstream effects of cytokine-producing capsule administration, the local and systemic levels of IL-10 and IL-12 were monitored over time via ELISA (fig. S5). The encapsulated cytokine-producing cells continued to produce detectable local levels of cytokines for up to 6 months when the study was concluded. Systemic levels of both cytokines were undetectable at 3- and 6-month timepoints, suggesting minimal cytokine leakage into systemic circulation. These comprehensive findings highlight the potential of locally produced cytokines to modulate the local immune response and effectively prevent initiation of fibrosis on biomaterial surfaces.

### IL-10 prevents fibrosis through the suppression of inflammatory response, while IL-12 prevents fibrosis by inhibiting the TGF-β pathway in intraperitoneal immune cells

Next, the underlying mechanism of cytokine-mediated fibrosis prevention was evaluated utilizing single-cell RNA (scRNA) sequencing. Briefly, 10 capsules (RPE, RPE-IL10 or RPE-IL12) were co-administered with empty microcapsules within the IP space of healthy C57BL/6J mice. Each RPE, RPE-IL10, and RPE-IL12 capsule contained 10,000 cells in a 1.5mm capsule, while empty microcapsules were fabricated at 0.5mm size. The capsules were retrieved at day 7 post-implantation, and the cells were collected from the capsule surface and surrounding IP fluid or the spleen. This approach allowed evaluation of both local and systematic effects induced by RPE-IL10 or RPE-IL12 in comparison to RPE control capsules upon introduction of foreign materials (in this case, empty microcapsules). Immune cell compositions, and expression signatures in each treatment group were analyzed and six distinct clusters using Uniform Manifold Approximation and Projection (UMAP) embedding were identified. Based on standard cell-type-specific markers, these clusters corresponded to granulocytes, monocytes (including macrophages), dendritic cells, B cells, T cells, and natural killer (NK) cells (Fig. 3A and fig. S6A). scRNA-seq analysis revealed that the composition of immune infiltrate in local space was significantly altered by cytokine-producing capsules (Fig. 3B). Compared to control RPEs, RPE-IL10 increased the intraperitoneal monocytes by 8.95% (*p* = 1.50e-12), while RPE-IL12 increased the intraperitoneal monocytes by 20.89% (*p* = 1.48e-64) (table S1). Notably, both cytokines significantly reduced the proportion of granulocytes, T cells, and NK cells present in the IP space by at least 3% (*p* < 1.1e-10 for all) and mildly altered (increased or decreased) the B cells and dendritic cell proportion by less than 3% (*p* < 0.05 for all) (table S1).

**Fig. 3.**
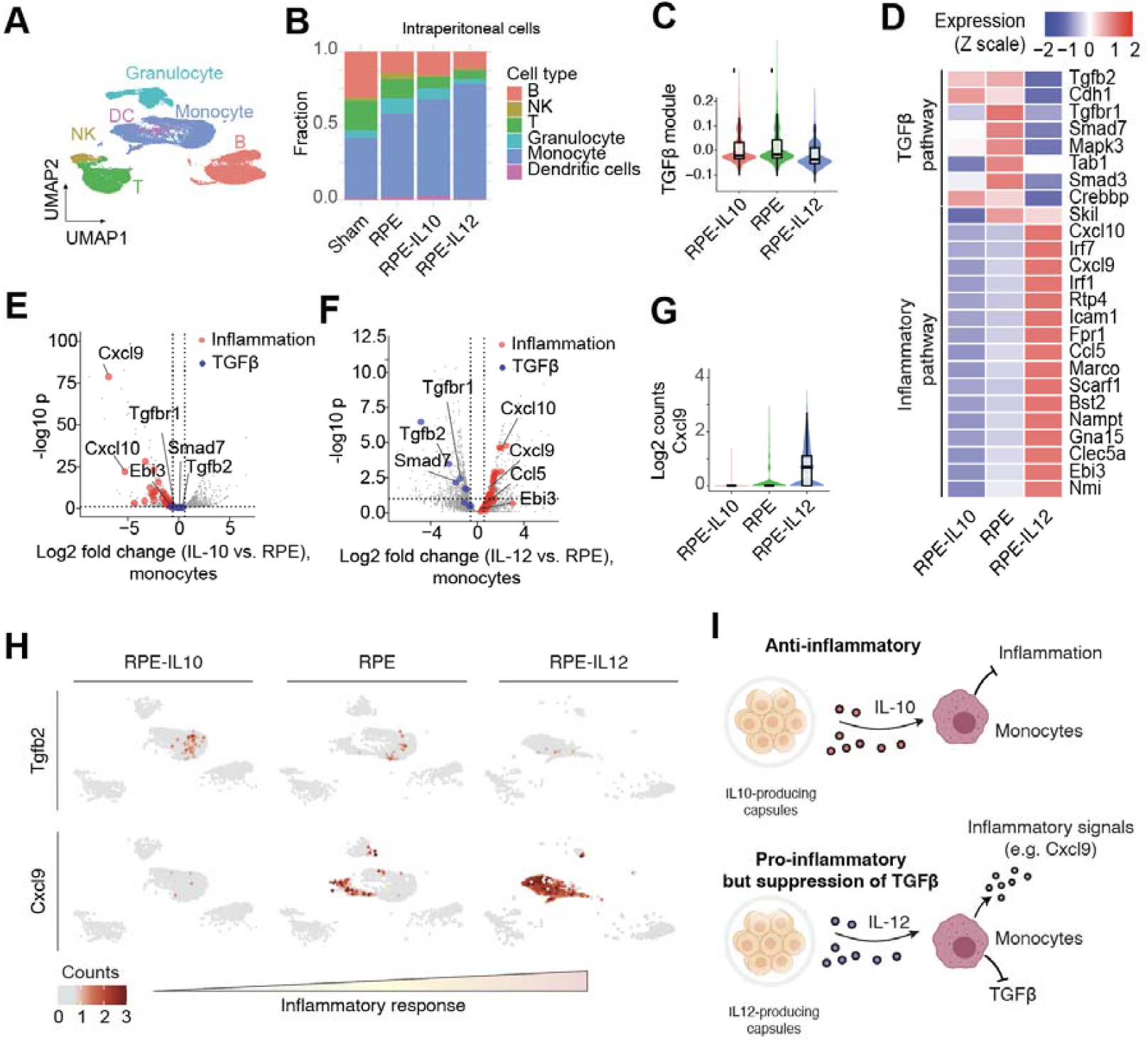
IL-10 prevents fibrosis by suppressing inflammation, while IL-12 prevents fibrosis by inhibiting the TGF-β pathway. (**A**) UMAP embedding of individual cells pooled from all samples. Resulting clusters are classified by immune cell type based on cell-specific expression profiles. (**B**) Composition of local immune cell identities recruited to eac implant site. (**C**) Violin plots showing the TGF-β module score in monocytes with RPE-IL10, RPE, and RPE-IL12. TGF-β module score was calculated based on expression levels of RPE-IL12-downregulated genes in the TGF-β pathway (p=2.9e-5 for RPE-IL10 vs. RPE, p=2.9e-16 for RPE-IL12 vs. RPE). (**D**) Heatmap showing z-scaled expression levels of representative genes from the inflammatory and TGF-β pathways. (**E** and **F**) Volcano plots showing the differential expression in monocytes treated with RPE-IL10 (E) and RPE-IL12 (F), compared to RPE alone. (**G**) Violin plot showing the inflammatory cytokine Cxcl9 in monocyte with RPE-IL10, RPE, and RPE-IL12 (p=2.9e-16 for both RPE-IL10 vs. RPE and RPE-IL12 vs. RPE). (**H**) Expression levels of Tgfb2 and Cxcl9 i intraperitoneal immune cells with RPE-IL10, RPE and RPE-IL12. (**I**) Schematics representing how RPE-IL10 and RPE-IL12 mediated fibrosis suppression. All p values are based on cell count.

The previous report (*38*) demonstrated that macrophages are indispensable to the fibrotic cascade in response to the implanted biomaterial. Despite that, our results showed that both anti-fibrotic RPE-IL10 and RPE-IL12 enriched the macrophage-containing monocyte population. To investigate the effect of RPE-IL10 and RPE-IL12 on fibrotic pathways, including inflammatory response and TGF-β (*39, 40*) pathway in monocytes, differential expression profiles of monocytes treated with each cytokine were compared to RPE alone. Here, the TGF-β pathway was investigated as it was shown to play a vital role in promoting fibrosis (*40*). Examining the expression of the TGF-β module indicated a significant decrease in TGF-β expression in the RPE-IL12 group compared to the RPE control (Normalized enrichment score (NES) = −2.1, *p* = 1.0e-3) (Fig. 3, C and D, and table S2). However, RPE-IL10 had mild but not statistically significant effects on the fibrotic TGF-β pathway (NES= −0.97, *p* = 0.49) (Fig. 3, C and D, and table S3). In analyzing the effects on the inflammatory pathway (Fig. 3D), it showed that RPE-IL10 significantly suppressed genes associated with the inflammatory response. In contrast, RPE-IL12 significantly promoted genes related to inflammatory response. Looking more closely, RPE-IL10 led to the downregulation of inflammatory Cxcl9 expression in monocytes (*p* < 2.2e-16) (Fig. 3E). Flow cytometry further confirmed an increased proportion of anti-inflammatory M2-like macrophages (CD11b□CD206□) and a reduced proportion of pro-inflammatory M1-like macrophages (CD11b□CD86□) in the RPE-IL10 group (fig. S7). The closer examination of RPE-IL12 revealed that it downregulated profibrotic Tgfb2 (*41*) (*p* = 4.9e-8) (Fig. 3F). As expected, in contrast to RPE-IL10, RPE-IL12 showed a higher expression of Cxcl9 (*p*=2.9e-16) (Fig. 3G). In addition, gene sets related to allograft rejection and antigen processing cross-presentation revealed that RPE-IL10 suppressed genes related to allograft rejection (NES = − 1.74, *p* = 2.8e-4) and antigen processing cross-presentation (NES = −1.74, *p* = 2.1e-3) (table S3). On the other hand, RPE-IL12 upregulated genes related to allograft rejection (NES = 1.32, *p* = 0.004) and antigen processing cross-presentation (NES = 1.72, *p* = 4.0e-6) (table S2). Expression of Tgfb2 was decreased in RPE-12 (top graphs), and expression of Cxcl9 was reduced in RPE-IL10 group (bottom graphs) (Fig. 3H). These results demonstrate that RPE-IL10 and RPE-IL12 exert distinct anti-fibrotic mechanisms; RPE-IL10 suppresses inflammation, while RPE-IL12 inhibits the TGF-β pathway (Fig. 3I).

Finally, the differences in local vs. systematic effects of cytokines on the gene expression profiles in the IP and spleen in RPE-IL10 or RPE-IL12 groups were compared to RPE group (fig. S6). RPE-IL10 does not significantly affect the gene expression in the spleen related to inflammatory gene sets, including allograft rejection (NES = −1.21, *p* = 0.13) and antigen processing cross-presentation (NES = −1.01, *p* = 0.43), indicating the effect of RPE-IL10 is primarily localized (fig. S6, B and C, and table S4). In contrast, RPE-IL12 significantly promoted inflammatory response, including allograft rejection (NES = 1.64, *p* = 7.8e-4) and antigen processing cross-presentation (NES = 2.04, *p* = 6.9e-6), and suppress TGF-β pathway (NES = −1.96, *p* = 5.8e-4) in splenic monocytes, indicating the effect of RPE-IL12 is systematic (fig. S6, B and C, and table S5). These results concluded that IL-10 is an ideal cytokine to prevent fibrosis with minimal systematic effect on the host’s immune cells.

### IL-10 producing cells prevent fibrosis of encapsulated xenogeneic islets, and enable long-term normal glycemic control in an immunocompetent diabetic mouse model

To evaluate the efficacy of RPE-IL10 and RPE-IL12 in protecting xenogeneic islets, we co-implanted cytokine-producing capsules in the IP space in streptozotocin (STZ)-induced diabetic C57BL/6J mice together with alginate-encapsulated human islets (2,000 islet equivalents; Fig. 4, A and B). RPE, RPE-IL10, or RPE-IL12 cells were encapsulated as described previously, and 10 capsules per mouse were co-administered with microcapsule. As in the fibrosis study, islet capsules were fabricated at 0.5 mm size to trigger a more robust FBR upon implantation. Following capsule administration, non-fasting blood glucose (BG) levels were drastically decreased and restored to normal glycemia levels in all groups. However, after 3 weeks, the RPE group failed glycemic correction (Fig. 4C), while the RPE-IL10 (Fig. 4D) and RPE-IL12 (Fig. 4E) groups demonstrated continued BG controls in diabetic mice up to 50 days with average BG levels below 250 mg/dL (which is commonly considered as healthy condition in the field).

**Fig. 4.**
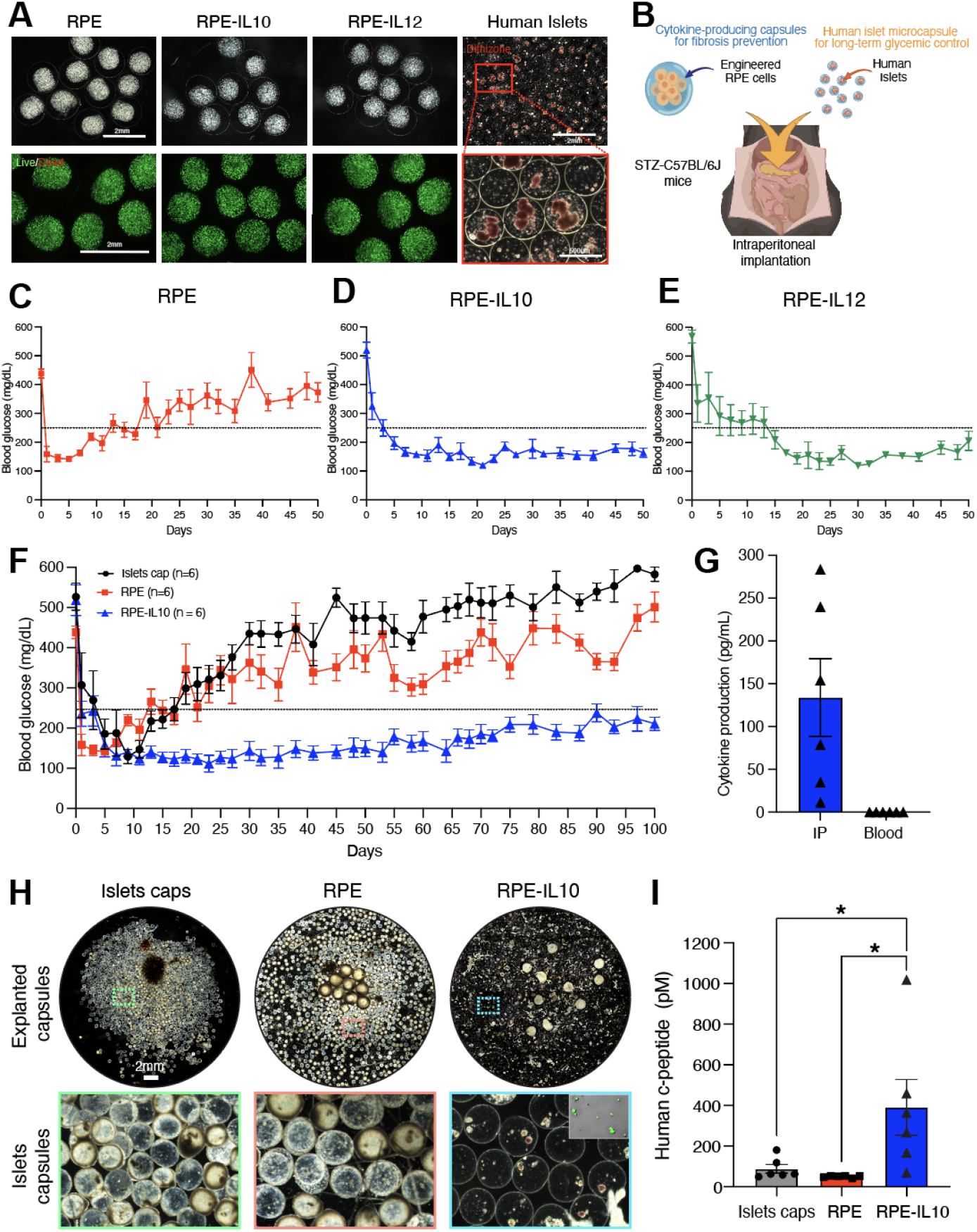
RPE-IL10 capsules co-delivered with encapsulated human islets achieved glucose correction in immunocompetent STZ-induced diabetic C57BL/6J mice. (**A**) Representative images of preimplant capsules. Dark-field images of alginate encapsulated 10,000 non-engineered RPE, RPE-IL10, and RPE-IL12 cells per capsule in a 1.5 mm diameter. After encapsulation, RPE, RPE-IL10, and RPE-IL12 demonstrate good viability. Live; green, dead; red. Dithizone staining (in red) shows β-cells inside the 0.5 mm diameter microcapsules. (**B**) Cytokine-producing RPE capsules are implanted with encapsulated human islets in the IP space of STZ-induced diabetic mice. (**C** to **E**) Blood BG levels of RPE (C), RPE-IL10 (D), and RPE-IL12 (E) groups delivered with encapsulated 2,000 IEQ islets up to 50 days (n = 6). Mice with BG levels below dashed grey lines (250 mg/dL) were considered to be in normoglycemic condition. (**F**) BG levels of RPE and RPE-IL10 capsules with islets microcapsules and islets microcapsules alone for 100 days (n = 6). (**G**) IL-10 local concentration (IP) and systemic concentration (blood) from the RPE-IL10 group at day 100. (**H**) Representative dark-field images of retrieved islets cap, RPE, and RPE-IL10 groups post 100 days implantation. Scale bar, 2mm. Inset in RPE-IL10; live/dead image of explanted islet capsules. (**I**) Human C-peptide levels in serum at day 100 post-transplantation for islets cap, RPE, and RPE-IL10 groups. All error bars denote mean□±□s.e.m of biological replicates. One-way ANOVA with Bonferroni correction was used for statistical analysis (*p < 0.033).

To access the durability of glycemic correction associated by cytokine-producing cells, BG levels of the RPE-IL10 group were monitored for 100 days in diabetic mice. RPE group co-administered with human islets (labeled as RPE) and only islets capsule group (labeled as islet cap) were used as controls. RPE-IL12 was not extended due to off-target complications (ascites) observed in a subset of animals, and subsequent analyses therefore focused on IL-10. RPE-IL10 with islet treatment demonstrated robust and extended BG regulation for 100 days compared to islet cap and RPE groups, which both failed before 3 weeks (Fig. 4F). Importantly, the sustained metabolic benefit in REP-IL10 group coincided with continued local cytokine availability, as IL-10 remained detectable in IP fluid at day 100 (Fig. 4G), indicating sustained secretion from the encapsulated cytokine-producing cells. Consistent with a cytokine-mediated attenuation of FBRs, explanted islet microcapsules from RPE-IL10 mice showed minimal pericapsular fibrotic overgrowth with highly viable islets relative to RPE and islets-only controls (Fig. 4H). Additionally, the concentration of human C-peptide, a surrogate biomarker for insulin production, was measured from the serum separated from mouse blood at day 100 post-transplantation. Significantly higher levels of C-peptide secretion were detected in the RPE-IL10 group compared with controls, supporting preservation of islet viability and function (Fig. 4I). Taken together, these findings support a coherent sequence in which persistent local IL-10 reduces fibrosis, preserves islet function, and sustains durable glycemic control, all achieved without systemic immunosuppression.

### IL-10-producing cells sustain local cytokine production and are well tolerated in a non-human primate

To evaluate clinical translatability, cynomolgus macaques received IP implantation of alginate-encapsulated cells in a pilot design: one animal with RPE-IL10 capsules (n = 1) and a control with non-engineered RPE capsules (n = 1). This study was not powered for efficacy and was intended to assess feasibility, pharmacokinetics, and tolerability. Previously published RPE-IL2 data (*14*) are referenced only for historical context. To assess cytokine leakage into the systemic circulation, cytokine levels in serum were measured. IL-10 levels showed transient systemic elevation on day 1 post-implantation, becoming undetectable by day 4 (Fig. 5A). In the RPE control animal, cytokines were undetectable at all time points. Similarly, data from our previous study with RPE-IL2 capsules demonstrated transient IL-2 detection in serum on day 3 but not by day 5 (*14*), consistent with brief, low-level systemic exposure early after implantation rather than sustained leakage. To assess local cytokine production, IP fluid was collected one-month post-implantation. In contrast to earlier findings with RPE-IL2 capsules (*14*), where cytokine levels were undetectable due to fibrotic encapsulation, IL-10 remained detectable in the RPE-IL10 group, suggesting sustained local cytokine production and reduced fibrosis (Fig. 5B). Pharmacodynamic effects of IL-10-producing cells were evaluated by monitoring CD4^+^CD25^high^ T_reg_ frequencies in peripheral blood. Flow cytometry of peripheral blood mononuclear cells (PBMCs), collected before implantation and after explantation, revealed T_reg_ expansion in the blood stream following RPE-IL10 implantation (Fig. 5C). General toxicity indicators, including body weight, body temperature, and platelet count, remained within normal ranges throughout the study (Table. 1). IL-10-related toxicities, particularly in the liver, kidneys, and lungs, were also assessed, as elevated IL-10 levels can suppress immune responses and impair organ function (*42-45*). Histopathological examination of these organs via H&E staining revealed no abnormalities, which aligns with the reported histology of healthy NHP organs (*46*) (fig. S8). Complete blood counts and blood chemistry showed no significant changes in red blood cell, white blood cell, or lymphocyte counts (table. S6). Kidney function was monitored through creatinine, blood urea nitrogen (BUN), and potassium levels (Table. 2). Although creatinine levels were slightly below baseline, this was unrelated to treatment, as levels were already low prior to implantation. BUN and potassium levels remained within healthy ranges. Liver function was assessed by measuring aspartate aminotransferase (AST), alanine aminotransferase (ALT), and gamma-glutamyl transferase (GGT) levels, and no significant changes were observed (Table. 3). These findings demonstrate that IL-10-producing cells support sustained, localized cytokine production without systemic exposure or toxicity, highlighting their safety and translational potential.

**Table 1.** General toxicity assessment: Longitudinal monitoring of body weight, body temperature, and platelet count.

|  | Day 0 | Day 1 | Day 4 | Day 7 | Day 14 | Day 21 | Day 28 | Normal Range |
| --- | --- | --- | --- | --- | --- | --- | --- | --- |
| Body Weight (kg) | 3.6 | 3.6 | 3.5 | 3.45 | 3.45 | 3.8 | 3.75 | N/A |
| Temperature (°C) | 34.3 | 36.9 | 37.3 | 37.5 | 37.1 | 37.7 | 37.6 | N/A |
| Platelet count (x10<sup>3</sup>/μL) | 427 | 349 | 191 | 546 | 529 | 586 | 628 | 154.56-636.2 |

**Table 2.**
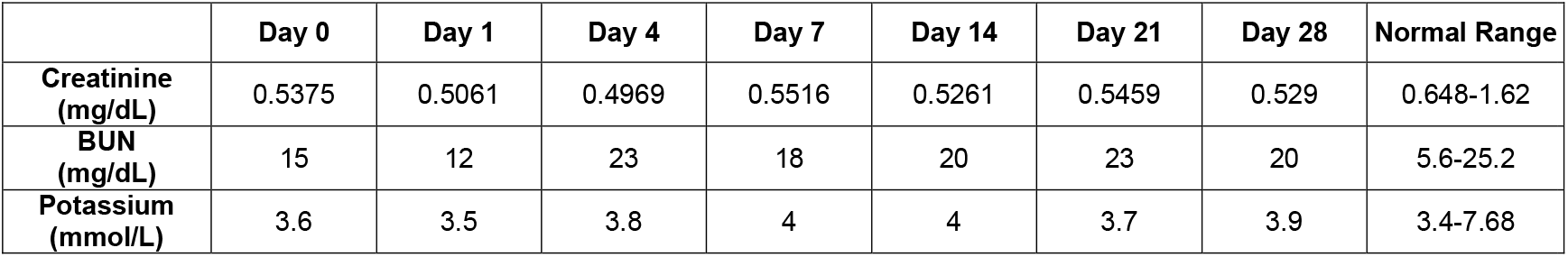
Kidney function assessment: Longitudinal analysis of serum creatinine, blood urea nitrogen (BUN), and potassium levels.

**Table 3.**
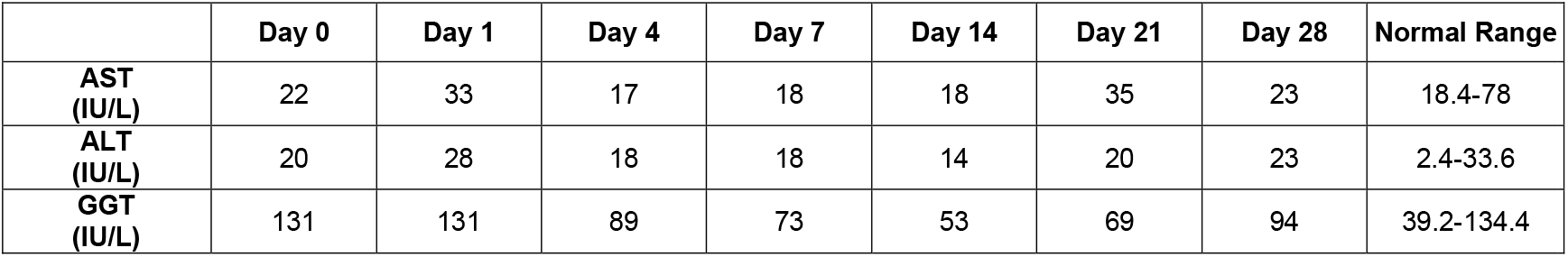
Liver function assessment: Longitudinal analysis of aspartate aminotransferase (AST), alanine aminotransferase (ALT), and gamma-glutamyl transferase (GGT) levels.

**Fig. 5.**
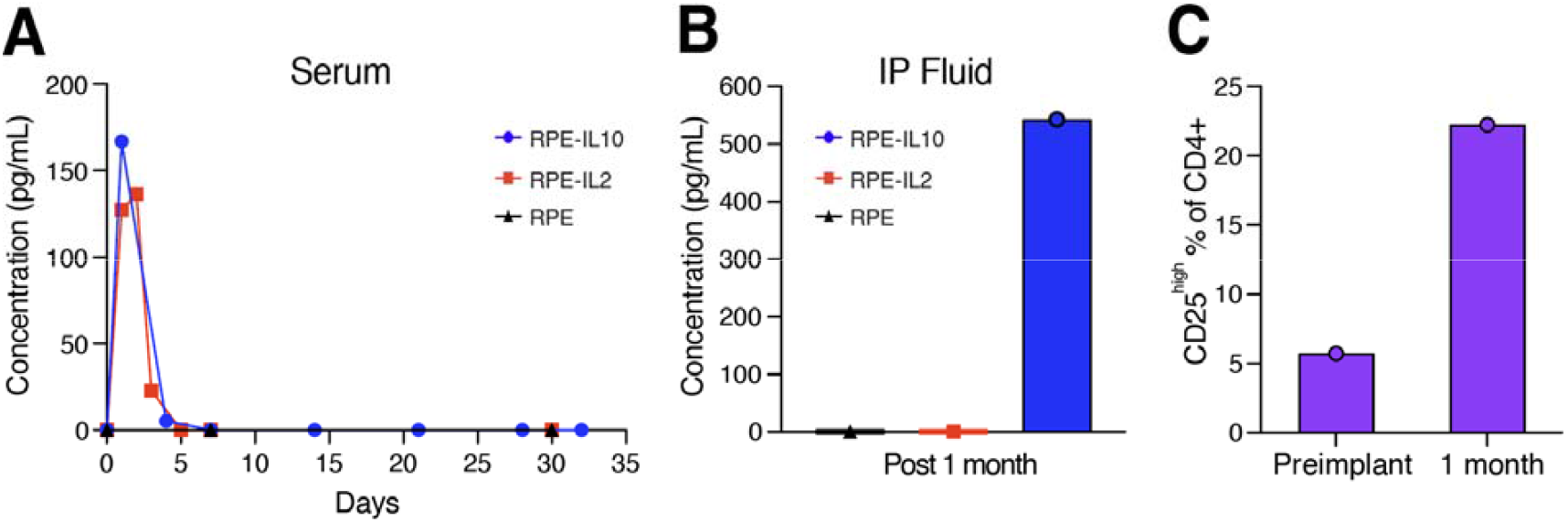
IL-10-producing cells showed sustained release of IL-10 and are well tolerated in a NHP. (**A**) Cytokine levels in the serum were measured over time via ELISA following administration of RPE-IL10, RPE-IL2, or control RPE capsules in each NHP. (**B**) Cytokine concentrations in the IP fluid were measured by ELISA one month after administration of RPE-IL10, RPE-IL2, or control RPE capsules in each NHP. (**C**) Changes in the frequency of CD4^+^CD25^high^ regulatory T cells in blood were assessed before capsule implantation and after explantation of RPE-IL10 capsules.

## DISCUSSION

This study presents a significant advancement in cell-based therapeutics by addressing one of the most pressing challenges: immune rejection and fibrosis of implanted cell therapies. By integrating a clinically translatable alginate-based platform with localized immunomodulatory cytokine delivery, our approach supports the robust and prolonged functionality of implanted cell therapeutics. Our findings build upon previous efforts to mitigate immune rejection in cell therapy platforms. Earlier strategies focused on chemical surface modifications, implant geometry alterations, or covalent polymer analogs to reduce immune responses and fibrosis (*9, 16, 23, 47*). While these approaches demonstrated promising preclinical results, translating them into clinical practice has proven challenging due to inconsistent outcomes and limited long-term efficacy. In parallel, clinical experiences with cytokine delivery by direct injection or slow-release depots have often been constrained by short half-lives, repeated dosing requirements, and systemic exposure-related toxicities (*34*). The delivery of IL-10 and IL-12 through our platform reduced fibrotic deposition in immunocompetent mouse models over six months, a substantial improvement compared to prior methods. Importantly, this effect was achieved without systemic toxicity, as cytokine secretion remained localized at the implantation site. These observations suggest that this cytokine-producing cell platform can reshape cytokine pharmacokinetics toward sustained, local exposure while minimizing systemic leakage, which may be difficult to achieve with bolus injections or conventional depots of cytokines.

The scRNA-seq analysis suggests a coherent mechanism for fibrosis mitigation. In the intraperitoneal compartment, IL-10 reprogrammed monocytes/macrophages by down-shifting inflammatory and antigen-presentation modules. This local tuning aligns with reduced pericapsular fibrosis and preserved islet function, without implying global immunosuppression. In contrast, IL-12 demonstrated a unique mechanism of action by modulating the TGF-β pathway, a critical mediator of fibrotic processes. Interestingly, IL-12 delivery led to localized inflammatory responses while simultaneously inhibiting fibrosis. This dual action highlights IL-12’s complex immunomodulatory role, as observed in our scRNA-sequencing analysis, which revealed the pleiotropic effects of TGF-β in promoting both anti-inflammatory signaling and fibrosis (*40, 41*). Despite its localized inflammation, IL-12’s ability to suppress extracellular matrix deposition positions it as a potential target for further investigation.

The inclusion of IL-10 and IL-12 into an alginate-based platform represents a key innovation in local immune modulation. Previous attempts to address graft rejection relied on systemic immunosuppressive therapies or engineered cell types such as cytokine-producing engineered β-cells (*37*) or immunomodulatory engineered MSCs (*48, 49*), which are associated with inconsistencies in cell survival, dose control, and spatial distribution. Other immunomodulatory approaches, such as checkpoint protein-releasing microgels (*50, 51*), demonstrated short-term efficacy but often required additional immunosuppressive regimens. In comparison, our platform offers several advantages. First, it enables sustained, localized cytokine delivery tailored to specific therapeutic needs. Second, it eliminates the requirement for systemic immunosuppression, thereby minimizing adverse immune-related effects. Third, the platform’s ability to achieve long-term efficacy in preventing xenogeneic graft rejection under stringent immune conditions highlights its translational robustness. Importantly, we observed concurrent local IL-10 detection, reduced pericapsular fibrosis, and preservation of human C-peptide with durable glycemic control in a diabetic mouse model, supporting a biologically linkage between cytokine delivery, fibrosis mitigation, and function. Collectively, these features distinguish our approach as a transformative advancement in cell delivery platforms.

The clinical implications of this work are particularly promising for T1D and other autoimmune diseases. Treatments aimed at slowing or reversing the fibrotic cascade have targeted modulating myofibroblast activation, reducing inflammation, inhibiting ECM deposition, reducing collagen synthesis, and/or enhancing ECM degradation, and many are currently being evaluated in clinical trials. Examples of anti-fibrotic treatments include drugs targeting i) the TGF-b pathway, such as Pirfenidone, which is an FDA-approved treatment for patients with idiopathic pulmonary fibrosis (*52*), ii) major growth factor pathways, such as Nintedanib, which is a receptor tyrosine kinase inhibitor (*52*), iii) the Pi3K pathway such as Parsaclisib which is a Pi3Kg inhibitor (*53*), iv) the JAK pathway such as Ruxolitinib which is JAK1/JAK2 inhibitor that is FDA-approved for patients with myelofibrosis (*54*), and v) glucose and lipid metabolism such as Semaglutide which is a GLP-1 receptor agonist that is FDA-approved for type 2 diabetes mellitus (*55*). Because many cell types and signaling pathways are implemented in fibrosis, there is no shortage of drug targets to attempt to ameliorate this process. Unfortunately, the side effects of these drugs can be severe and thus remain a major hurdle for drug development. Safer treatments for patients are urgently needed.

IL-10 has been extensively studied for its role in immune modulation, with previous reports demonstrating its ability to suppress insulitis and delay diabetes onset in animal models (*56, 57*). In humans, elevated IL-10 levels have been associated with improved β-cell protection and immune tolerance (*57*). However, prior attempts to deliver IL-10 using viral vectors or recombinant proteins faced challenges, including immunogenicity, toxicity, and rapid cytokine degradation (*58*). Our platform overcomes these limitations by providing long-term, localized IL-10 delivery, ensuring sustained therapeutic effects while minimizing systemic exposure. Notably, therapeutic colocalization of every cytokine-producing capsule with islet capsules is not required in the IP cavity; the peritoneum functions as a permissive diffusion compartment in which a relatively small fraction of cytokine-producing capsules can maintain a bioactive cytokine condition that influences the broader graft environment. Mechanistically, IL-10-associated reductions in fibrosis are consistent with decreased expression of pro-fibrotic markers and preserved islet function observed at late time points.

Beyond T1D, this local immunomodulatory platform holds significant potential for other autoimmune, inflammatory, and infectious diseases. For instance, the ability to deliver IL-10 locally could be applied to conditions such as rheumatoid arthritis, inflammatory bowel disease, and organ transplantation, where immune modulation is critical for disease management (*59-62*). The platform’s modularity also allows for the incorporation of other cytokines or therapeutic agents, expanding its utility across various medical applications. Future studies exploring these opportunities will further establish the platform as a versatile and clinically translatable solution for fibrosis-related disorders. To assess clinical translatability, we conducted a pilot NHP evaluation focused on feasibility, pharmacokinetics, and tolerability rather than efficacy. In the NHP study, the cytokine-producing cells successfully prevented fibrosis and immune rejection of cell therapeutics, mirroring the results observed in mouse models. Notably, the localized delivery of IL-10 maintained cytokine concentrations at the implant site without detectable systemic toxicity post one month implantation, demonstrating its safety in a clinically relevant model.

While this study demonstrates considerable advancements, certain limitations must be addressed in future research. Although IL-10 delivery effectively prevented fibrosis and graft rejection in immunocompetent mouse and NHP models, the systemic effects of IL-12 observed in splenic monocytes raise concerns regarding its long-term safety and immunogenicity. We note that fluid accumulation was not observed with the IL-12 dose in healthy mice, suggesting disease-context interactions in diabetic models; nevertheless, dose titration and temporal control will be required for safe translation. Future work will include refined IL-12 dosing with mechanistic validation, and safety features for IL-10 delivery such as inducible expression or kill-switches to allow on-demand cessation. Additionally, while the platform’s success has been demonstrated in mouse and NHP models, further investigation into disease-specific fibrotic models, such as idiopathic pulmonary fibrosis and liver fibrosis, will help expand its therapeutic applications. Long-term surveillance for infection susceptibility or tumor immunity under sustained IL-10 exposure will also be incorporated into translational studies.

In conclusion, this study establishes a clinically translatable platform that prevents fibrosis and immune rejection of cell-based therapeutics while offering broad potential for addressing fibrotic diseases. By combining a clinically translatable alginate platform with localized cytokine delivery, we provide a robust and durable solution to fibrosis and immune rejection in cell therapies. IL-10’s ability to prevent fibrosis and preserve graft functionality underscores its potential as a therapeutic candidate for T1D and other autoimmune diseases. The platform’s capacity to achieve sustained, localized cytokine delivery with minimal systemic toxicity ensures both safety and efficacy, marking a significant advancement over existing immunomodulatory strategies. NHP feasibility and tolerability further support clinical potential. Overall, this work represents a paradigm shift in the prevention and treatment of fibrosis, offering new opportunities for advancing cell therapies, regenerative medicine, and fibrosis-focused therapeutics.

## MATERIALS AND METHODS

### Study design

The objective of this study was to develop and evaluate a localized immunomodulatory platform that uses encapsulated, cytokine-producing cells to prevent immune-mediated rejection and fibrosis in cell-based therapies. We hypothesized that sustained, local delivery of immunoregulatory cytokines, such as IL-10, from alginate-encapsulated engineered RPE cells would attenuate host immune responses and preserve graft. The study included in vitro assessments of cytokine secretion and cell viability, followed by in vivo evaluation in murine and non-human primate models. For rodent studies, engineered cells were co-delivered with human islets and implanted into STZ-induced diabetic C57BL/6J mice to assess fibrosis, glycemic control, and cell function over time. For translational relevance, the platform was further validated in cynomolgus macaques, where encapsulated IL-10-secreting cells were laparoscopically delivered to the peritoneal cavity, and local and systemic immune responses were monitored over time. Capsules were retrieved at various time points for analysis of fibrosis, cell viability, cytokine levels, and immune infiltration using ELISA, live/dead imaging, scRNA-seq, and immunohistochemistry. Additional assays included C-peptide measurements, RT-qPCR, and flow cytometry. Experimental groups were randomly assigned where applicable. Detailed animal numbers, sample sizes, and data points for each experiment are provided in the corresponding figure legends. All animal protocols were approved by the Institutional Animal Care and Use Committees (IACUC) at Rice University and the University of Illinois Chicago, and were conducted in accordance with institutional guidelines.

### Cell Culture and Transfection

Cell culture and transfection procedures were conducted using materials from Fisher Scientific and Invitrogen, including cell culture media and associated reagents. Expression vectors and helper plasmids were obtained from VectorBuilder. Transfection reagents Lipofectamine 3000 and selection antibiotic (puromycin) were purchased from Invitrogen. The Opti-MEM media for transfection was purchased from Thermo Fisher Scientific. The ARPE-19 cell line, procured from ATCC, underwent regular testing for mycoplasma contamination, yielding negative results. Cells were cultured in Dulbecco’s Modified Eagle Medium (DMEM/F-12) supplemented with 10 % FBS and 1 % antibiotic-antimycotic (AA), with media changes occurring three times weekly. ARPE-19 cells were engineered to express cytokines of interest, following the established transfection protocols (*14*). Briefly, ARPE-19 cells were seeded into six-well plates at a cell density of 500,000 cells per well. The cells in the plate were left overnight in the incubator and primed with 2 mL of Opti-MEM serum-free media for 15 minutes before transfection. A 1:2 ratio of helper plasmid to plasmid expressing cytokine of interest was used to transfect cells. All cell lines were transformed according to the manufacturer’s protocol. After incubation of cells with transfection agents for 4 hours at 37 °C, the transfection medium was replaced with fresh culture media, and cells were left to incubate overnight. Then, the cells were selected for expression with puromycin for 2 weeks and expanded to quantify the expression via ELISA.

### Engineered Cell Encapsulation

For engineered cell encapsulation, SLG20 alginate (UP-LVG, NovaMatrix, Norway) was dissolved at 1.4 % w/v in 0.8% saline, followed by sterile filtration. Before encapsulation, engineered cells underwent trypsinization and centrifugation at 250 g for 5 minutes. Cell pellets were washed twice with Ca-free Krebs buffer and then resuspended in the alginate solution at a density of 5 x 10^6^ cells/mL (∼10,000 cells per capsule). Capsules were produced using a custom-built, two-fluid co-axial electrostatic spraying device (Harvard Apparatus), as described previously (*9, 14*). Alginate droplets were ejected from an 18 G co-axial needle (Rame-Hart) into a barium chloride crosslinking solution, forming hydrogel capsules with a diameter of 1.5 mm. The flow rate of both syringe pumps for core and shell solutions was adjusted to 5 to 6 mL/hr. Capsule size was controlled by adjusting the voltage to between 5.5 and 6 kV (Gamma High Voltage). The capsules were incubated in the crosslinking solution for 15 minutes, washed with HEPES buffer, and maintained using standard cell culture techniques.

### Human Islet Encapsulation

Human islets (Prodo Labs) were cultured in PIM(S) media from the same source. After centrifugation and washing with Ca-free Krebs buffer, the islets were resuspended in a 1.4 % SLG20 solution at a density of 20,000 islet equivalents (IEQ) per 2.7 mL. The islet-containing solution was loaded into a syringe with a 25 G blunt-tipped needle. Following the same fabrication procedure as 1.5 mm capsules, microcapsules were adjusted to a size of 0.5 mm by setting the voltage to 10 kV with a flow rate of 200 μL/min. After washing, 400 μL aliquots of microcapsules with 2,000 IEQ cells were prepared for implantation. Additionally, empty microcapsules, without cells, were fabricated under the same conditions for use in a fibrosis assay.

### Creation of STZ-Induced Diabetic Model

For in vivo studies, a mix of male and female C57BL/6J mice (Charles River Laboratories) aged 8 to 10 weeks was employed. All animal studies were approved by Rice University’s Institutional Animal Care and Use Committee (IACUC). To induce insulin-dependent diabetes, healthy C57BL/6J mice received STZ treatment. STZ solution at a concentration of 7.5 mg/ml (50 mg/kg of STZ) was injected into the intraperitoneal (IP) space for five consecutive days. BG levels and weights of the mice were measured after a one-hour fasting. Only mice with BG levels exceeding 350 mg/dL for two consecutive days were deemed diabetic and selected for islet transplantation.

### Intraperitoneal Surgical Implantation of Capsules in Mice Models

Immunocompetent C57BL/6J mice (8 weeks old) were weighed, anesthetized with 1 to 4 % isoflurane in oxygen on a heating pad, and administered subcutaneous Ethiqa XR based on weight. After shaving and sterilizing their abdomens with betadine and isopropanol, a 0.5 to 1 cm midline incision was made through the skin. The peritoneal wall was grasped with forceps, and a 5 mm incision was made along the linea alba. For the fibrosis study (1-, 3-, and 6-month time points), 10 capsules containing engineered cells in a 1.5 mm diameter size were implanted into the IP cavity, along with 0.4 mL of empty microcapsules (0.5 mm size). For the scRNA-seq study, 10 capsules containing engineered cells and 0.4 mL of empty microcapsules were implanted into the IP space for 7 days. For the islet study with STZ-induced diabetic mice, portions of empty microcapsules were replaced with islet microcapsules (with 2,000 IEQ per mouse). The incision was sealed with a suture.

### NHP Capsule Implantation

Adult Mauritian cynomolgus monkeys were used for capsule implantation studies. All procedures were approved by the University of Illinois Chicago (UIC) IACUC and adhered to the Guidelines for the Care and Use of Laboratory Animals. Under general anesthesia and sterile conditions, a 2-cm supraumbilical incision was made, and a 5-to 12-mm trocar was inserted to establish pneumoperitoneum with CO□ (10-14 mmHg). A laparoscopic camera was used for visualization, and a 2-cm incision was made for a second trocar to deliver capsules. Capsules, resuspended in 50 mL of saline, were administered via a catheter-tip syringe and distributed throughout the IP cavity. Incisions were closed in layers using Vicryl sutures (3-0 for muscle, 4-0 for skin). After capsule implantation, the health of the NHPs was closely monitored through regular assessments, including body temperature and weight measurements, as well as comprehensive bloodwork.

### Blood Glucose Monitoring

Blood glucose levels were monitored three times weekly after the transplantation of islet capsules, conducted without fasting. Mice displaying BG levels below 250 mg/dL were categorized as normoglycemic. Monitoring persisted until all mice reverted to a hyperglycemic state, at which juncture they were euthanized, and the capsules were recovered.

### Retrieval of Capsules

At a designated point in the implantation period, mice from each study were euthanized using cardiac puncture procedures followed by cervical dislocation. An incision along the abdomen skin and peritoneal wall was made using forceps and scissors. IP fluid and capsules within the cavity were collected using 10 mL of sterile PBS. Separation of all capsules from the IP fluid was carried out, followed by several washes with Krebs buffer. The explanted capsules were prepared for further imaging and promptly snap-frozen for future analysis.

### Capsule Imaging and Cell Viability Assay

The capsules were gently washed with Krebs buffer and transferred to 35 mm Petri dishes for bright- and dark-field imaging using an EVOS microscope. Under 2X magnification, images were acquired and stitched to observe the entire dish. Fluorescent imaging of cells stained with the live/dead assay (Invitrogen, catalog # L3224) was performed to assess encapsulated cell viability in both pre- and post-implant capsules. Five capsules from each group underwent washing with PBS and staining with 2 μM calcein AM and 4 μM EthD-1 in complete media. Following a 30-minute incubation, capsules were imaged using an EVOS microscope with fluorescence filters. Live cells were visualized with a GFP filter in green, while dead cells were observed with a Texas-Red filter in red.

### scRNA-seq Analysis

On day 7 post-implantation, animals were euthanized, and cell populations from IP fluids and retrieved capsules were collected to analyze the local effects of the therapeutic. Spleens from each group were also collected to examine systemic immune cell populations. After removing red blood cells, cells were resuspended in 1 mL of media for cell counting. Propidium Iodide was added to stain dead cells following the manufacturer’s instructions. Live cells, sorted at 1000 cells/μL density in DMEM containing 10 % FBS, underwent next-generation sequencing at Baylor College of Medicine. Throughout sample processing and cell sorting, cells were maintained on ice. The Single Cell 5’ Gene Expression Library was generated using the Chromium NextGEM Single Cell Immune Profiling Solution 5’v2 protocol by 10x Genomics. The libraries were sequenced on an Illumina NovaSeq 6000 flow cell. Transcripts in each cell were counted using the 10x Cell Ranger 5.0.1 pipeline, with genome mapping to the mm10 genome build using STAR v2.7.2a. The scRNA-seq data was visualized using UMAP embedding in Loupe Browser 5.0 to identify immune cell populations. At this stage, six cell populations were readily apparent. The resulting clusters were assessed for the expression of common immune cell marker genes and were then classified as specific immune cell types based on their expression profiles. The cell identities of each of the six clusters were resolved using the following markers: CD3E (T cells), PRF1 (NK cells), CD19 (B cells), FN1 (monocytes and macrophages), ITGAX (dendritic cells), and LY6G and HDC (granulocytes). These markers assigned individual cell barcodes to their corresponding immune cell type in the Loupe browser. Cell proportions were calculated between cytokines for each immune cell type to assess changes in the infiltrated immune composition. The significance of these changes was calculated using Fisher’s exact test in R (v3.6.1). Differential expression in monocytes between different treatments was derived from the Loupe browser. TGF-β module score was calculated based on expression levels of RPE-IL12-downregulated genes (Tgfb2, Cdh1, Tgfbr1, Smad7, Mapk3, Tab1, Smad3, Crebbp, Skil) in BIOCARTA_TGFB_PATHWAY using addModule method in Seurat (v3.1.5). Expression levels of representative genes from the BIOCARTA_TGFB_PATHWAY pathway and HALLMARK_INFLAMMATORY_RESPONSE were z-scaled and plotted in a heatmap using ComplexHeatmap (v2.8.0).

### Enzyme-Linked Immunosorbent Assay (ELISA)

For preimplant capsules, a single capsule was added to a 96-well plate (n = 8) after encapsulation in 200 µL of culture media at 37 °C in a 5 % CO_2_-humidified atmosphere. Capsule supernatant was collected after 24 hours. For explant capsules, a single capsule from each mouse post-retrieval was added to a 96-well plate (n = 6) in 200 μL of media for 24 hours, and the plate was kept in the incubator. Capsule supernatant was collected from each well and assayed. Capsule supernatants were assayed at 10x, 100x, and 1000x dilutions depending on the ELISA kit. For the local and systemic concentration of cytokines, IP fluid was assayed at 1x and 10x dilution, and blood was assayed at 2x and 10x dilution. ELISAs were obtained commercially for mIL10 (R&D Systems, Catalog #: M1000B), mIL12 (R&D Systems,Catalog #: M1270), mIL2 (R&D Systems, Catalog #: M2000). The assay was run according to the manufacturer’s protocols. All the samples were run in duplicates.

### Human C-Peptide Assay

Following manufacturer protocol, blood from each mouse post-explant was assayed in duplicate without dilution (ALPCO, Catalog#: 80-CPTHU-E01.1). All the standards were also assessed in duplicates.

### Immunofluorescence Staining for Confocal Imaging

Retrieved microcapsules were washed with Krebs buffer, fixed in 4% paraformaldehyde overnight at 4 °C, and permeabilized with 1 % Triton X-100. Following blocking with a 1 % bovine serum albumin (BSA) solution, samples were incubated with antibody cocktails (diluted at 1:200 Alexa Fluor 488 anti-mouse CD68 Antibody (Cat# 137012, BioLegend), 1:200 Anti-mouse α-Smooth Muscle-Cy3 (Cat# C6198, Sigma-Aldrich), and DAPI (Cat# R37606, Invitrogen) in 1 % BSA) for 1 hour at room temperature. After washing, samples were transferred for imaging using a Nikon A1-Rsi confocal microscope.

### RT-qPCR Analysis

Total RNA was extracted from 100 μL of retrieved microcapsules (empty capsules with a 0.5 mm size) using the RNeasy Mini Kit (Cat# 74104, Qiagen). The extracted RNA was converted to cDNA for RT-qPCR using the high-capacity cDNA reverse transcription kit (Cat# 4368814, Applied Biosystems). Real-time qPCR (Bio-Rad) was performed using SYBR Green (Cat# A25742, Applied Biosystems), and reactions were run in triplicates under specified conditions. Data analysis was conducted using the 2^□ΔΔCT^ method, comparing relative RNA levels after normalization to mouse ActB and Empty Cap control. The primer details are listed in table. S7.

### Flow cytometry

All antibodies were commercially sourced, prepared fresh on the day of staining, and stored in the dark at 4°C or on ice to preserve their efficacy. Cells were prepared for staining for flow cytometry as previously reported (*14*). Samples were washed with 200 µl of cell staining buffer (Cat# 420201, Biolegend) prior to staining. For NHP samples, cells were stained with anti-human CD4 (Cat#555349, BD, 1:200) and anti-human CD25(Cat# 560920, BD, 1:100) for 20 minutes on ice in the dark. After staining, samples were washed three times with cell staining buffer and fixed for an hour at room temperature with fixation buffer (Cat# 420801, Biolegend). After fixation, cells in each sample were resuspended in cell staining buffer and run through a 40-μm filter before acquisition on a Sony MA900. Single-color, FMO controls, as well as a negative unstained control, were prepared for each analysis. Gating method was described in the fig. S9.

For mouse samples, cells were stained with viability dye (Cat# 423114, Biolegend, 1:1000), anti-mouse CD11b (101224, Biolegend, 1:50), anti-mouse CD206 (Cat #141716, Biolegend, 1:40), and anti-mouse CD86 (Cat #105014, Biolegend, 1:20) for 30 minutes on ice in the dark. Then, cells were washed with fluorescence-activated cell sorting (FACS) buffer and fixed with fixation buffer (Cat# 88-8824-00, Thermo Fisher) for 30 minutes at room temperature. After fixation, cells in each sample were resuspended in 300 μl of FACS buffer and run through a 40-μm filter for eventual analysis using an SA3800 Spectral Analyzer. Single-color, unstrained control and FMOs were included. The gating strategy is shown in the fig. S10.

### Histology

The Baylor Pathology and Histology Core facilitated the processing, sectioning, and histological analysis of NHP organs. Samples were paraffin-embedded, excess paraffin was meticulously trimmed, and sagittal sections were prepared and stained with hematoxylin and eosin (H&E) for detailed examination.

### Statistical Analysis

All statistical analyses were conducted with GraphPad Prism 9. One-way or two-way ANOVA with Bonferroni multiple-comparison correction was used to determine p values (*\*\*\*\*P < 0*.*0001, ***P < 0*.*0002, **P < 0*.*002, *P < 0*.*033*).

## Supplementary Materials

**The PDF file includes:**

Figs. S1 to S10

Table S1

Legends for tables S2 to S5

Tables S2 to S5 (Merged excel file)

Tables S6 and S7

## Acknowledgments

We thank Baylor College of Medicine’s Genomic and RNA Profiling Core and Human Tissue Acquisition and Pathology Core. We would also like to acknowledge the use of equipment at the Shared Equipment Authority of Rice University. We also thank Rice University animal resource facility staff for their assistance with animal research. All the schematics were made using Biorender.

## Author contributions

Conceptualization: B.K., A.M.N., D.V., and O.V. Methodology: B.K., D.V., A.M.N., Y.C., S.F., S.D., D.M., P.D.R., I.J., H.N., D.L., J.O., C.H., and O.V. Investigation: B.K., D.V., A.M.N., Y.C., S.F., S.D., D.M., P.D.R., I.J., H.N., D.L., M.S.G, C.H., J.O., C.H., and O.V. Visualization: B.K., D.V., Y.C., and O.V. Project administration: B.K., A.M.N., D.V., P.D.R., C.H., and O.V. Supervision: O.V. Writing—Original draft: B.K., A.M.N., D.V., and O.V. Writing—Review and editing: B.K., A.M.N., D.V., and O.V.

## Data and materials availability

All data are available in the main text or the supplementary materials.

