## Supplemental Figures for "Localized immunomodulation with cytokine-producing cells to mitigate host immune rejection responses in rodents and a non-human primate"

Boram Kim *et al*.

**The PDF file includes:**

Figs. S1 to S10
Table S1
Legends for tables S2 to S5
Tables S6 and S7

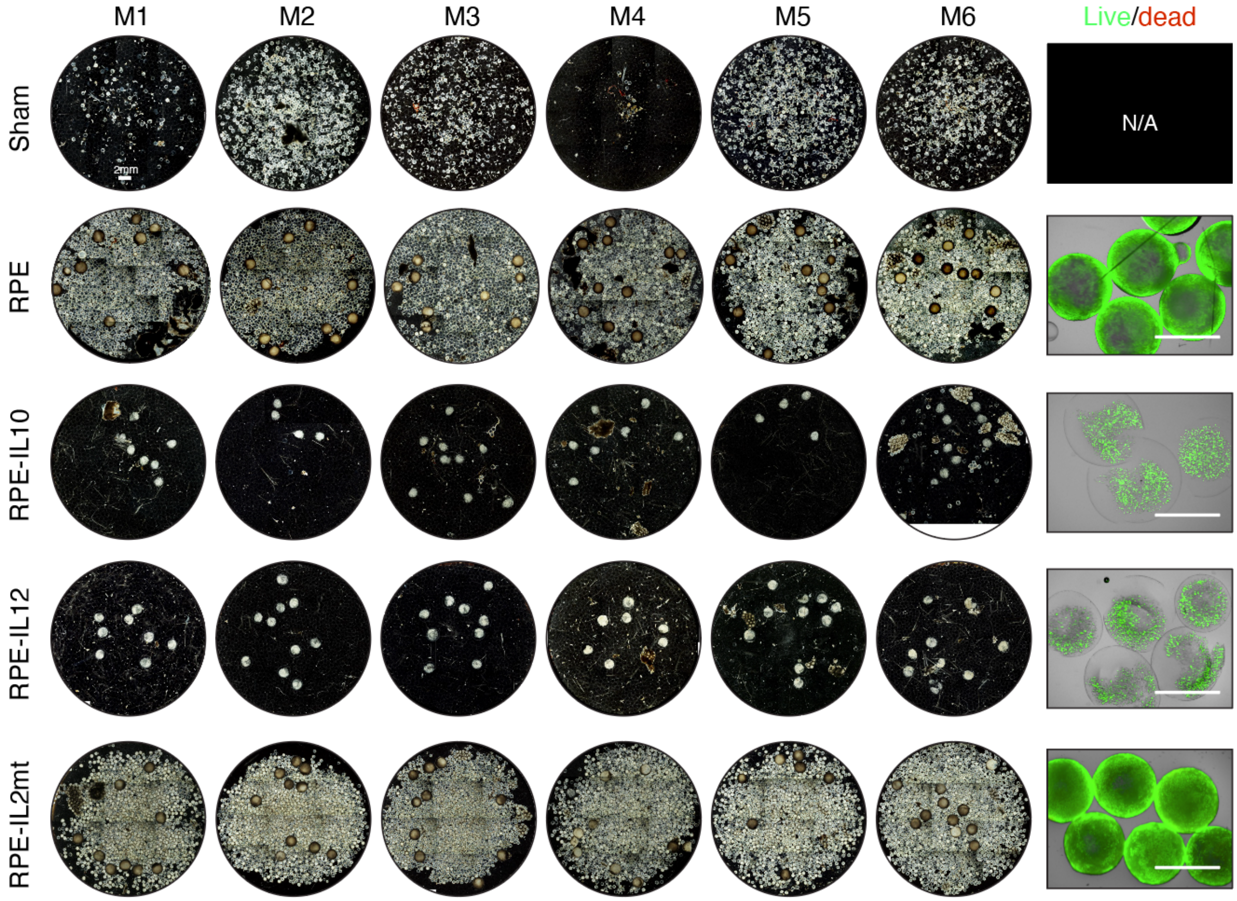


**Fig. S1. Short-term effect of cytokine-producing capsules in preventing fibrosis.** Dark-field images of explanted capsules from individual mouse post one month IP implants. The last column shows the representative live/dead staining images of explanted RPE capsules (Live, green; Dead, red). Scale bar, 2mm.

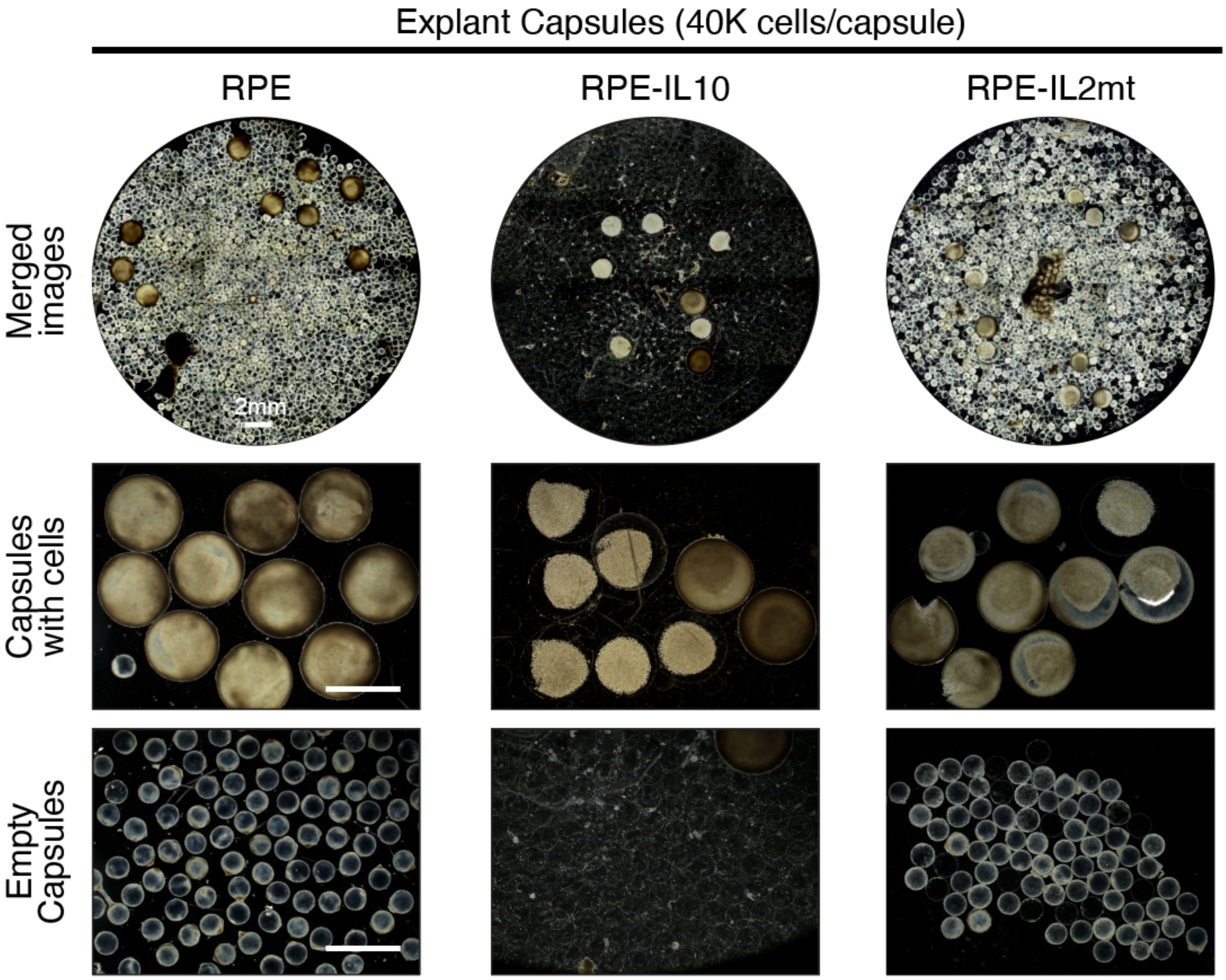


**Fig. S2. Short-term effect of high-dose cytokine-producing capsules on fibrosis.** Capsules were fabricated at 40,000 cells per capsule (high-dose condition). Representative dark-field images of explanted capsules post one month IP implants. Scale bar, 2mm.

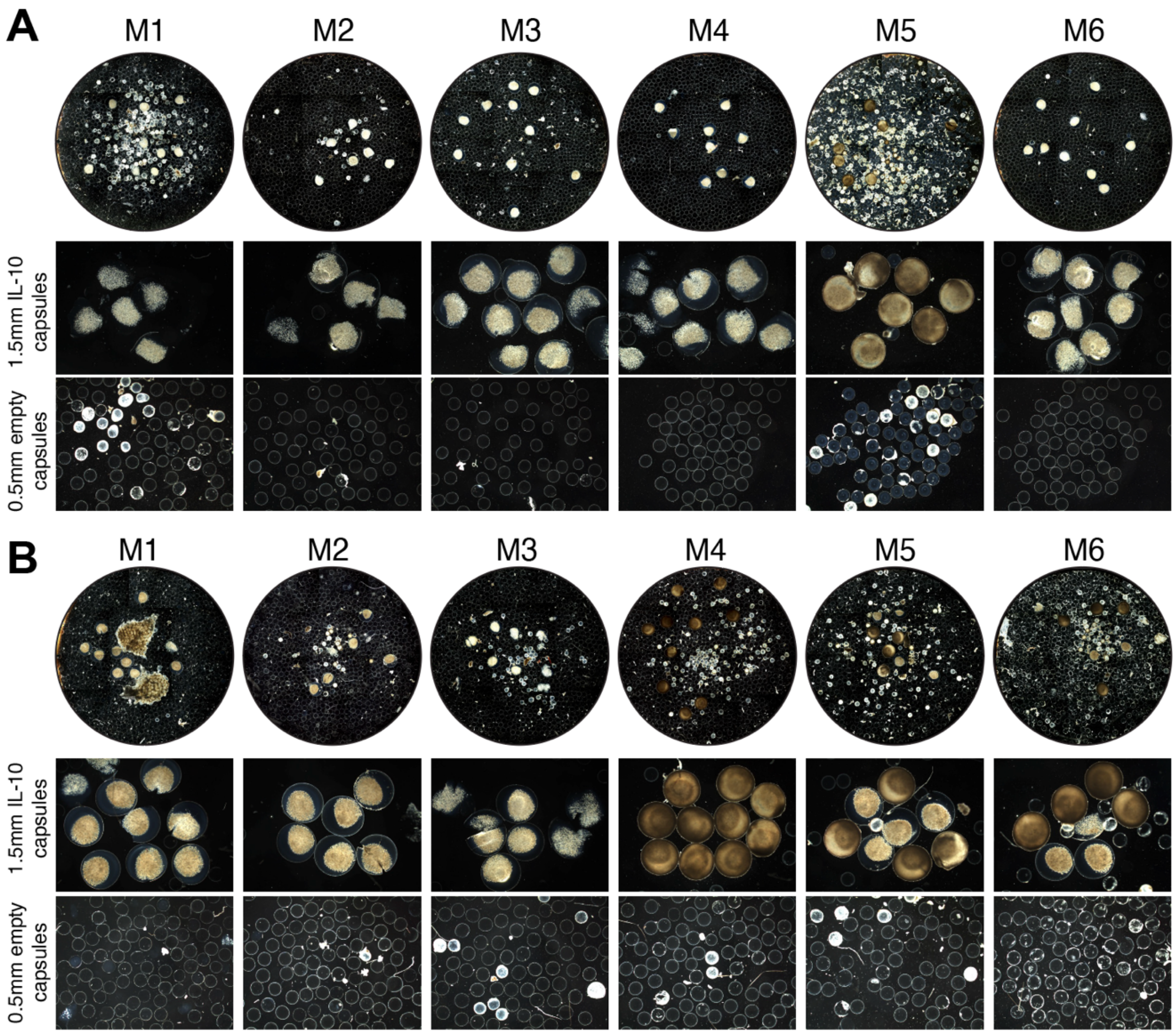


**Fig. S3. Long-term effect of RPE-IL10 in preventing fibrosis.** (**A** and **B**) Dark-field images of retrieved capsules after three months (A) and six months (B) implantation.

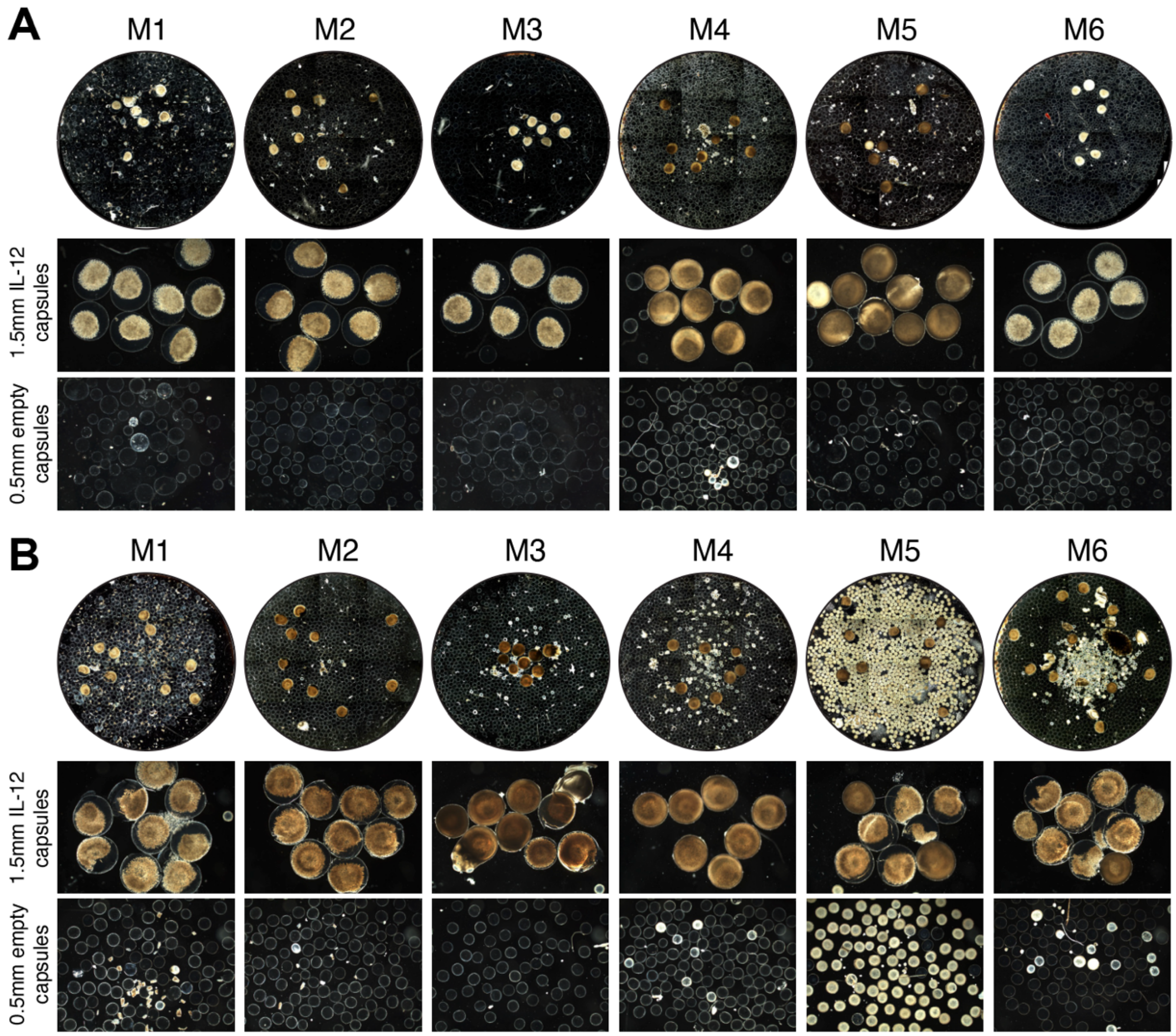


**Fig. S4. Long-term effect of RPE-IL12 in preventing fibrosis.** (**A** and **B**) Dark-field images of retrieved capsules after three months (A) and six months (B) implantation.

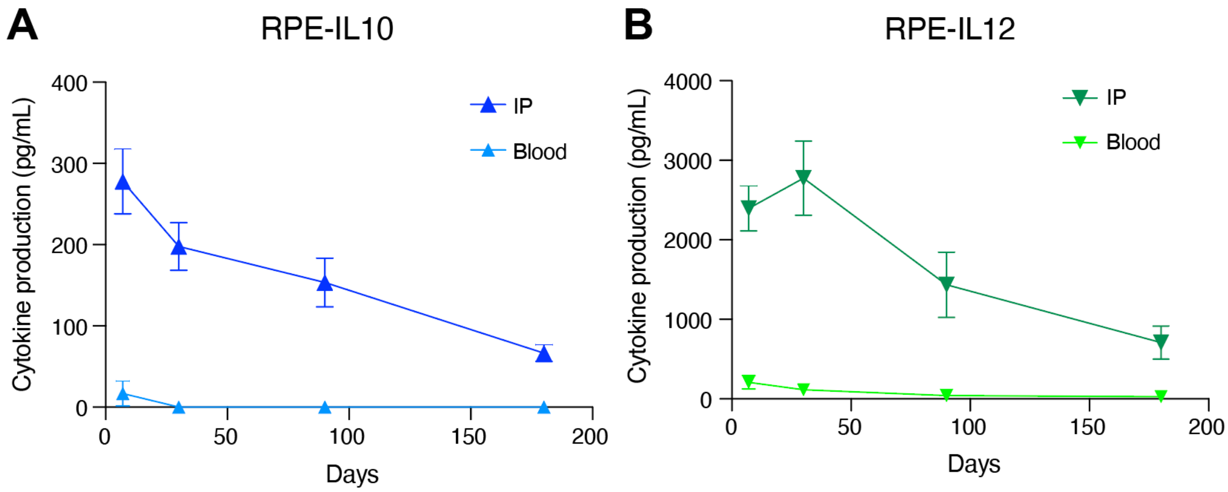


**Fig. S5. Cytokine concentrations in IP fluid vs bloodstream over time.** (**A**) RPE-IL10 implant group. (**B**) RPE-IL12 implant group.

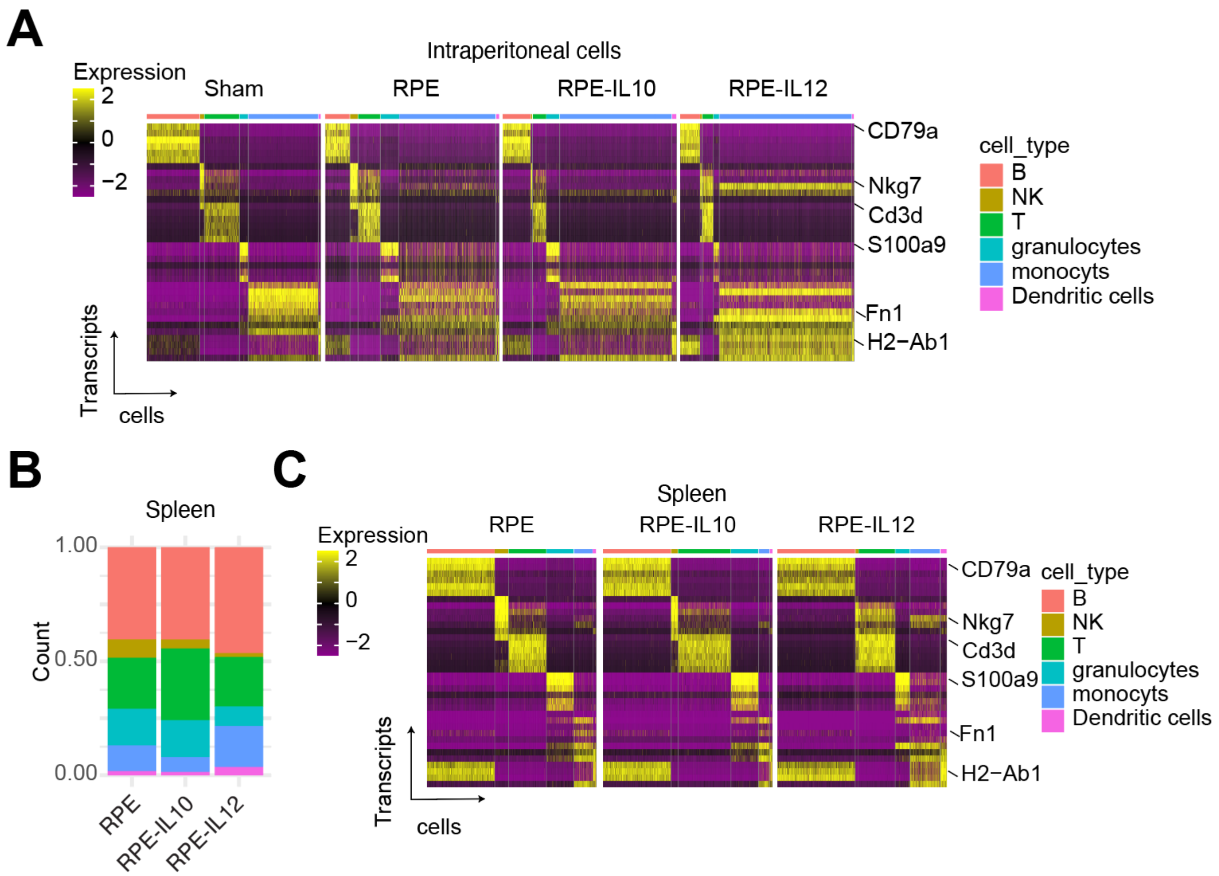


**Fig. S6. scRNA-seq analysis.** (**A**) Single-cell transcriptional profiles of markers used to define each cell immune cell type in intraperitoneal cells. (**B**) Composition of splenic immune cell identities. (**C**) Single-cell transcriptional profiles of markers used to define each cell immune cell type in splenic cells.

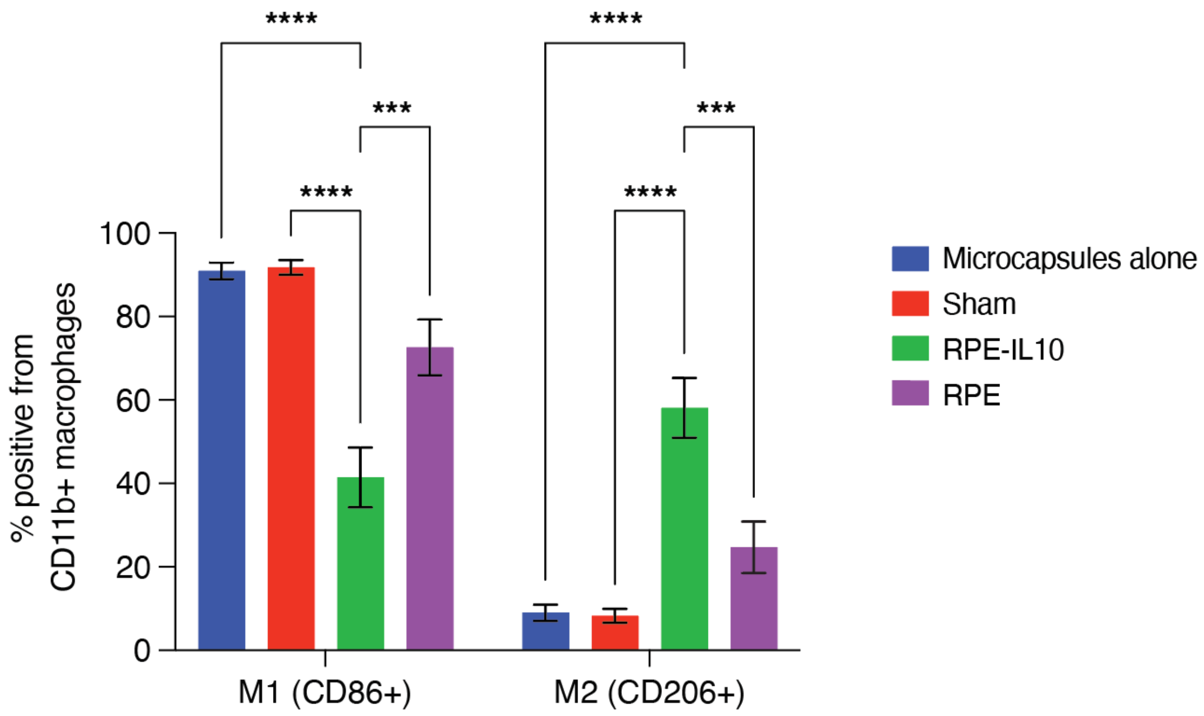


**Fig. S7. Flow cytometry on intraperitoneal cells of healthy mice at Day 7 post-implantation.** The frequency of M1-like macrophages (CD11b+CD86+) versus M2-like macrophages (CD11b+CD206+) was evaluated in the intraperitoneal space among the groups. All error bars denote mean ± s.e.m of biological replicates (n=5).

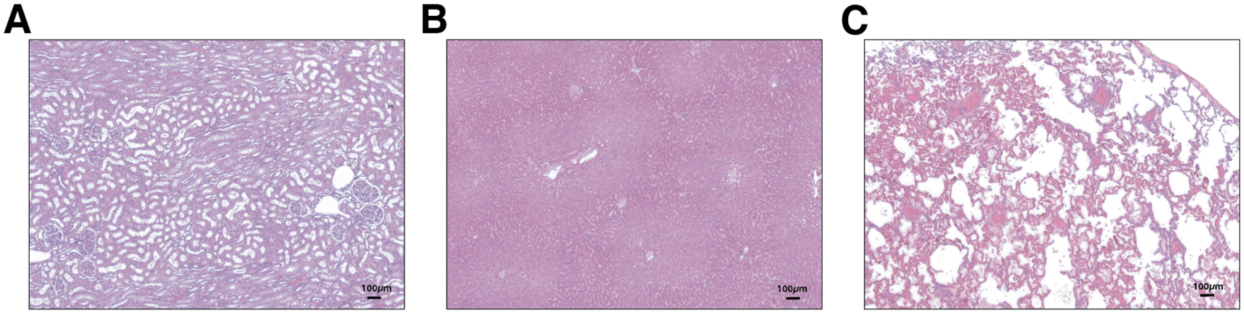


**Fig. S8. NHP histology images.** (**A** to **C**) Representative histology images of NHP kidney (A)**,** liver (B)**,** and lung (C) taken at 20x magnification. No notable clinical abnormalities were detected.

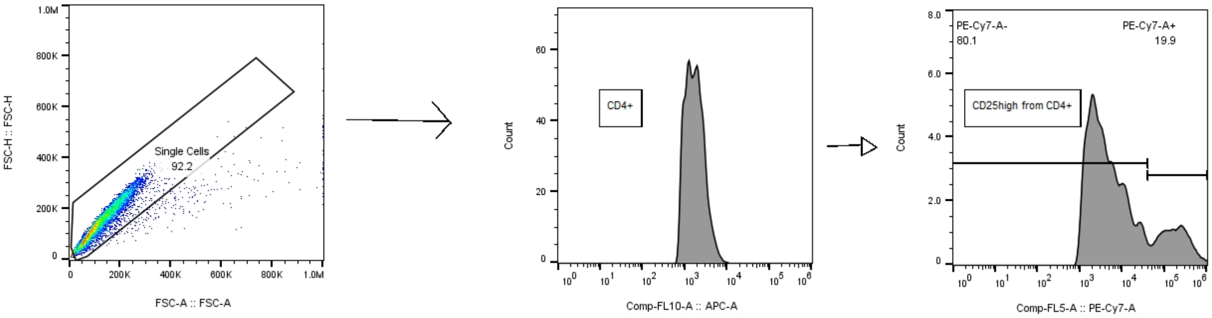


**Fig. S9. NHP flow cytometry gating strategy.**

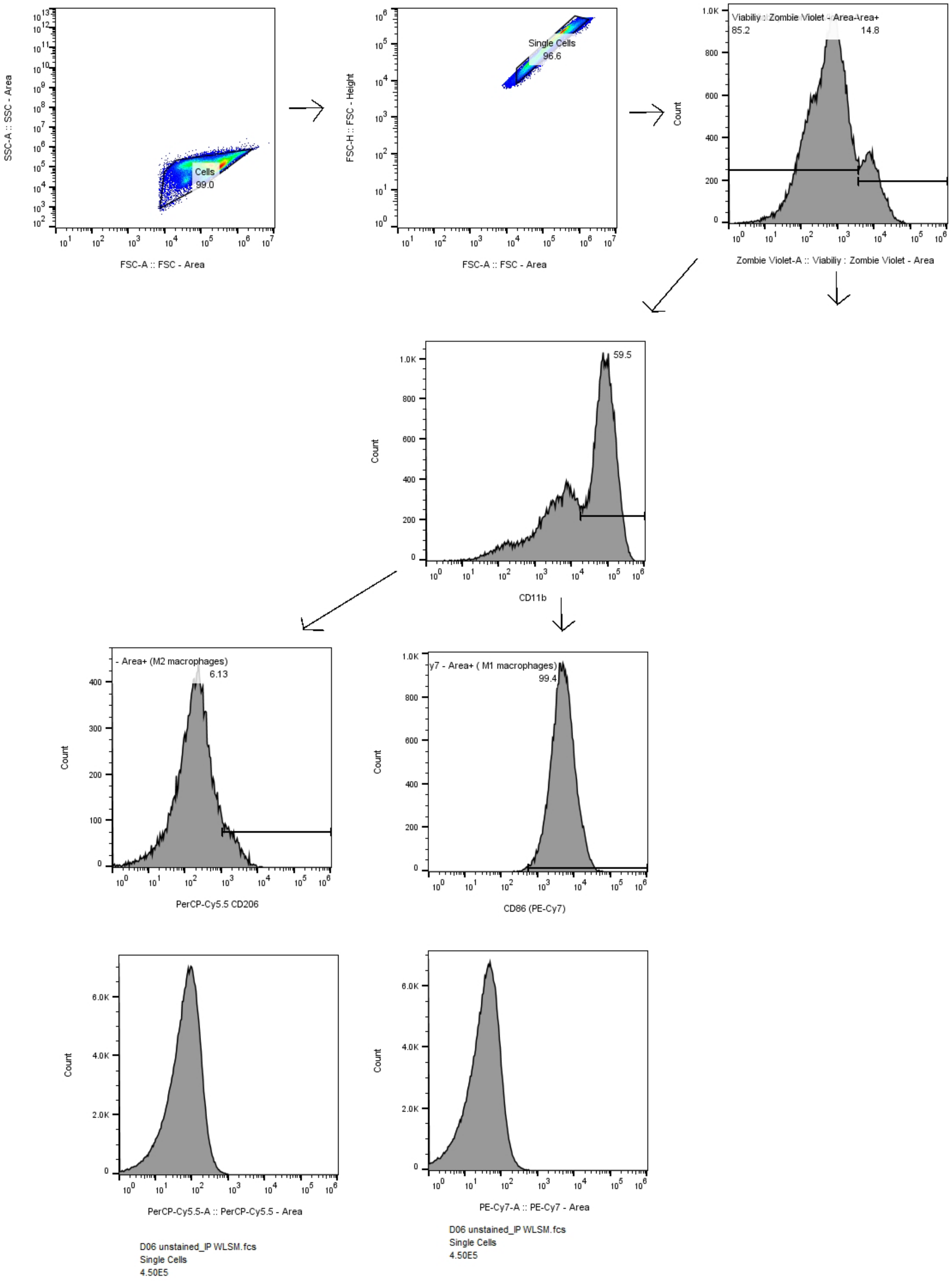


**Fig. S10. Flow cytometry gating strategy on mice samples.**

### Table S1. Composition of immune infiltrate in local vs. systemic space.

| Local: Percentage of immune cells in IP/capsule (%) | | | | | | | | | | | | |
|---|---|---|---|---|---|---|---|---|---|---|---|---|
| **Cell type** | **B** | **Other** | **DC** | **Other** | **NK** | **Other** | **T** | **Other** | **Granulo** | **Others** | **Mono** | **Other** |
| RPE | 14.3 | 85.7 | 1.8 | 98.2 | 4.5 | 95.5 | 12.8 | 87.2 | 10.6 | 89.4 | 56.1 | 43.9 |
| IL-10 | 16.2 | 83.8 | 2.5 | 97.5 | 0.8 | 99.2 | 7.8 | 92.2 | 7.6 | 92.4 | 65.1 | 34.9 |
| RPE | 14.3 | 85.7 | 1.8 | 98.2 | 4.5 | 95.5 | 12.8 | 87.2 | 10.6 | 89.4 | 56.1 | 43.9 |
| IL-12 | 11.6 | 88.4 | 1.1 | 98.9 | 1 | 99 | 6.2 | 93.8 | 3.2 | 96.8 | 77.0 | 23.0 |
| **Systemic: Percentage of immune cells in spleen (%)** | | | | | | | | | | | | |
| **Cell type** | **B** | **Other** | **DC** | **Other** | **NK** | **Other** | **T** | **Other** | **Granulo** | **Others** | **Mono** | **Other** |
| RPE | 40.5 | 59.5 | 1.9 | 98.1 | 8.1 | 91.9 | 22.4 | 77.6 | 16.1 | 83.9 | 11.1 | 88.9 |
| IL-10 | 40.5 | 59.5 | 1.5 | 98.5 | 3.9 | 96.1 | 31.5 | 68.5 | 16.2 | 83.8 | 6.4 | 93.6 |
| RPE | 40.5 | 59.5 | 1.9 | 98.1 | 8.1 | 91.9 | 22.4 | 77.6 | 16.1 | 83.9 | 11.1 | 88.9 |
| IL-12 | 46.5 | 53.5 | 3.7 | 96.3 | 1.7 | 98.3 | 21.7 | 78.3 | 8.6 | 91.4 | 17.8 | 82.2 |

* B, B cells; DC dendritic cells; NK, NK cells; T, T cells; Granulo, Granulocyte; Mono, monocytes.

**Table S2. Gene set enrichment analysis of differential analysis between RPE-IL12 vs. RPE in intraperitoneal monocytes.**

**Table S3. Gene set enrichment analysis of differential analysis between RPE-IL10 vs. RPE in intraperitoneal monocytes.**

**Table S4. Gene set enrichment analysis of differential analysis between RPE-IL10 vs. RPE in splenic monocytes.**

**Table S5. Gene set enrichment analysis of differential analysis between RPE-IL12 vs. RPE in splenic monocytes.**

*Note: Tables S2-S5 are provided as one merged .xlsx file.*

#### Table S6. NHP hematological data for the primates given RPE-IL10 capsules.

| Units / Days | RBCs | WBCs | Lymphocytes count | Platelet count |
|---|---|---|---|---|
| | (x10^6/μL) | (x10^3/μL) | (x10^3/μL) | (x10^3/μL) |
| Day 0 | 6.69 | 8.35 | 73.7 | 427 |
| Day 1 | 6.33 | 10.06 | 35.3 | 349 |
| Day 4 | 5.15 | 13.57 | 44.4 | 191 |
| Day 7 | 4.15 | 12.52 | 48.8 | 546 |
| Day 14 | 3.29 | 12.83 | 61.3 | 529 |
| Day 21 | 4.85 | 12.49 | 49.1 | 586 |
| Day 28 | 5.89 | 7.8 | 58.4 | 628 |

#### Table S7. List of RT-qPCR primers.

| Target | Primers (5'→3') |
|---|---|
| Mouse Alpha Smooth muscle actin (αSMA) | Forward: CGCTTCCGCTGCCCAGAGACT |
| | Reverse: TATAGGTGGTTTCGTGGATGCCCGCT |
| Mouse Collagen 1a1 (Col1a1) | Forward: CATGTTCAGCTTTGTGGACCT |
| | Reverse: GCAGCTGACTTCAGGGATGT |
| Mouse Collagen 1a2 (Col1a2) | Forward: GCAGGTTCACCTACTCTGTCCT |
| | Reverse: CTTGCCCCATTCATTTGTCT |
| Mouse β-actin (ActB) | Forward: GCTTCTTTGCAGCTCCTTCGTT |
| | Reverse: CGGAGCCGTTGTCGACGACC |
